# Factors regulating translation coupled mRNA degradation in yeast

**DOI:** 10.1101/2022.04.01.486758

**Authors:** Sudipto Basu, Raktim Maiti, Suman Hait, Ashesh Ray Chaudhuri, Sudip Kundu

**Affiliations:** Department of Biophysics, Molecular Biology and Bioinformatics, University of Calcutta, Kolkata, India; GenML Solutions(OPC) Pvt. Limited, Kolkata, India

**Keywords:** mRNA half-life, Co-translational degradation, Codon optimaity, mRNA secondary stucture, Ribosomal footprinting

## Abstract

Codon optimality is a major factor governing translation speed and efficiency, mRNA expression, and stability. Presence of optimal codons maintains a steady translation time, while non-optimal codons can lead to slow peptidyl transfer, prolonged translation time and frequent halt. Owing to this, there are co-translational surveillance protein complexes which monitor this process meticulously and can trigger the release of the mRNA transcript after prolonged halt. The surveillance protein complexes are closely associated with various degradation enzymes which can break down these released transcripts. Degradation of the released transcripts will be ensured if they contain amenable regions for enzyme binding and action. Here, we have used sequence data to estimate the relative abundance of non-optimal codons within a transcript. We found that transcripts with high relative abundance of non-optimal codon show reduced stability and half-life compared to transcripts with lower relative abundance of non-optimal codon. We have further integrated the relative abundance of non-optimal codons with internal unstructured segments for the transcripts, to show that the former is responsible for transcript release and the latter provides susceptible platform for endonuclease cleavage, which together is capable of destabilising and reducing the transcript’s half-life. Since this is a translational system, sequestration of mRNA into a dense forest of ribosomes is seen to prolong the transcript’s half-life even in the presence of destabilising factors.

## 1. Introduction

The life-cycle of an mRNA starts from a nascent RNA transcript being synthesized in the nucleus, undergoes various transcriptional post-processing checkpoints like capping, splicing, and polyadenylation, ultimately getting exported to the cytoplasm as mature mRNA. Once in the cytoplasm, the sole purpose of mature mRNA is to undergo translation for protein synthesis. During this process, there are protein complexes screening for the fidelity and efficiency of these mRNA transcripts. Transcripts are immediately triggered for degradation, if their fidelity or efficiency is compromised. The degradation of mature mRNA gives them a characteristic half-life. Cytoplasmic mRNA undergoes degradation via 5’-3’ or 3’-5’ degradation pathway after decapping and poly-A tail removal. In the case of exonucleolytic degradation, it is established that both 5’-3’ or 3’-5’ degradation depends on the terminal unstructured length of 5’ and 3’ respectively, while endonucleolytic cleavage depends on the presence of Internal Unstructured Segments (*IUS*) [1].

Another major gateway to redirect mature cytosolic mRNA to degradation is via co-translation surveillance system [2]. Co-translational surveillance screens for mRNA that trigger degradation including (i) encountering a premature stop codon [3], (ii) lacking a stop codon [4], and (iii) with stalled ribosomes [5]. The Ccr4-Not (carbon catabolite repressor 4-negative on TATA) is a conserved protein complex, working as co-translational surveillance system, mediating the transfer of transcripts to mRNA degradation machineries [6, 7]. On stalling, Ccr4-Not protein complex can initiate 3’-tail deadenylation [7], and subsequent activation of 5’-decapping holoenzyme (Dcp1-Dcp2) [8], making the transcript susceptible to both 5’-3’ or 3’-5’ mediated degradation [9]. A specific subunit Not5 (NTD domain) bridges an association between ribosome and Ccr4-Not complex [10] screening for flawed translation initiation, slow peptidyl transfer and decoding during elongation. During elongation, Not5 can sense a prolonged vacancy in the A-site resulting in a state where an *aa*_*tRNA*_ (amino-acyl tRNA) is occupied only at the P-site 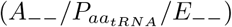. Such a situation often arises due to the presence of non-optimal codon in the transcript, for whom the respective *aa*_*tRNA*_ pool in the system is not so enriched. The NTD of Not5 on apprehending such a scenario communicates with the degradation machinery for immediate action [10]. As understood, codon optimality is one of the reasons which regulates the action of Not5 of Ccr4-Not complex, hereafter regulating mRNA degradation and half-life 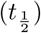.

Different mRNA transcripts show differential codon content. Over the years, this versatility has been linked to translational dynamics, protein expression, and abundance [11, 12, 13, 14]. It has been demonstrated earlier in *Saccharomyces cerevisiae* [15], *Schizosaccharomyces pombe* [16], *Escherichia coli* [17], zebrafish [18, 19] and in human and mammalian cell lines [20, 21] that codon optimality is a major determinant of mRNA stability. Moreover these studies are all limited in establishing a relation between non-optimal codon (*NOC*) content and mRNA half-life 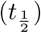. The non-optimal codons within the transcripts may be differentially distributed having various stretch lengths and frequencies, which can exert various effects on mRNA stability, but this correlation has never been explicitly explored. Therefore, we have introduced a metric which calculates the relative strength of non-optimal codons over optimal codons within an mRNA transcript to understand their effect on their respective half-lives.

Association of RNA Binding proteins stabilize mRNA transcripts by burying regions which are amenable for nuclease capture for degradation [1]. Ribosome, being a macromolecular ribonucleoprotein complex, plays a key role in the translation system. During translation, ribosomes function in clusters placed at equidistant regions on the transcript as polysomes. The density of ribosomes stationed on the transcripts plays a key role in stabilising the transcripts [22].

Here, we have integrated (i) multiple yeast genome wide mRNA half-life 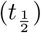 datasets, (ii) mRNA sequence data, (iii) mRNA secondary structure data and (iv) ribosomal footprinting data to understand the dynamics of translation linked mRNA degradation and stability. In this study, the (i) mRNA sequence information is used to extract derived parameters like Codon content index (*CI*) and Cumulative score(*cs*) and the (ii) mRNA secondary structure information is utilized to derive Internal Unstructured Segment (*IUS*) to address the issue. We found that *NOC* stretch whether they are present in large numbers or in larger lengths reduces translation speed and stalling occurs due to their reduced *aa*_*tRNA*_ concentration in the cellular pool. This increases the chances of releasing the transcripts for degradation and reduced stability. *IUS* within the released transcript provides suitable regions for endonuclease attachments and influences the mRNA stability. Despite the presence of *NOC* and *IUS* within a transcript, we have observed that polysomal association elevates mRNA stability and increases its half-life.

## 2. Materials and methods

### 2.1 Data collection

#### 2.1.1 *S. cerevisiae* transcript half-life and sequence data

Three non-redundant experimental half-life datasets of *S. cerevisiae* [23, 15, 24], are used for this study (Data S1). The annotated transcript sequence of *S. cerevisiae* is collected from Saccharomyces Genome Database [25].

#### 2.1.2 Ribosomal foot-printing Data

Ribosomal footprinting data provide a quantitative measurement of the number of ribosomes stationed on an mRNA transcript. It is generated using deep sequencing of the ribosomal protected fragments [26]. For this study, we have obtained the ribosomal footprinting data from GEO database (GEO ID: GSE75897) [27] (Data S1). This contains three sets of footprinting data which are extracted using three different techniques. They are Unselected, Dynabeads, and Ribominus.

#### 2.1.3 Internal Unstructured Data

PARS give the secondary structure data of mRNA at single nucleotide resolution, with the aid of nucleotide digestion and high-throughput sequencing [28]. Using PARS data, we have generated Internal Unstructured Segment(*IUS*), a derived parameter calculated using a window of a 12 nucleotides long moving from 5’ to 3’ end of the transcript with 6 nucleotides slide at a time, similar to the previously established method [1] (Data S1). The IUS (*ξ*) score is calculated as 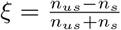. Here, *n*_*s*_ and *n*_*us*_ denotes the number of unstructured and structured segment in the transcripts. *ξ* is calculated for different threshold percentage. For example, in a window of 12 nucleotides, if more than 60% of the nucleotides can form base pairs (PARS score *>* 0), and less than 40% nucleotides are in an unpaired state (PARS score *≤* 0), we denote it as *ξ*_*s*60*us*40_. In a similar way, different percentage threshold is used for counting both structured and unstructured segments and calculate *ξ* score for each transcript.

### 2.2 Transcript Analysis

#### 2.2.1 Codon Content Index

Here, Codon Content Index (*CI*) is calculated as the normalized difference between the non-optimal codon (*NOC*) and the optimal codon (*OC*). *CI* value ranges from −1 to +1, where negative and positive *CI* for a transcript indicates the abundance of optimal and non-optimal codon respectively within that transcript (Data S2). The calculations are performed using in-house Python scripts. The mathematical representation of *CI* is as follow:

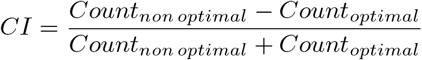

#### 2.2.2 Cumulative scoring for *NOC*

An mRNA transcript can contain multiple stretches of non-optimal codon of different lengths. We have estimated the frequency (*f*) of *NOC* stretches (*≥* 3) length for each transcript. Ribosomal stalling largely depends on the A-site and P-site, during incorporation of *aa*_*tRNA*_ and peptidyl transfer, and is independent of E-site. So, a stretch of more than two consecutive *NOC* could be responsible in keeping the process stalled for a considerable amount of time. So, we have introduced a combined scoring parameter (*cs*) for encompassing the effects of both length (*j*) and frequencies (*f*_*j*_) of these *NOC* stretches in a transcript (Data S2). *cs* can be mathematically represented as:

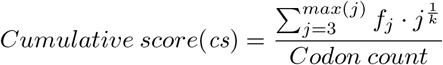

Here, *j* denotes the length of *NOC* stretch with a minimum value of 3. So, *cs* is actually a summation of all the *NOC* stretch lengths put to the power of *k* which is used as a weightage factor, and multipled by their respective *NOC* length frequencies(*f*) for a given transcript. The weightage factor (*k*) is varied from 0.01 to 10. For *k* = 1, the *cs*-score indicates the *NOC* content normalized by the total codon. The weightage factor is varied in such a way, such that for (i) *k <* 1, it elevates the effect of *NOC* stretch length (*j*) over frequency (*f*) on *cs*-score and (ii)*k >* 1, it elevates the effect of frequency (*f*) over *NOC* stretch length (*j*) on *cs*-score. The calculations are performed using in-house Python scripts.

### 2.3 Statistical analysis

All the statistical tests provided in this manuscript are performed using PAST v3.0 [29].

## 3. Results and Discussions

### 3.1 Non-Optimal Codon content regulates mRNA half-life

There are a handful of previous literature reporting the effect of *NOC* on mRNA half-life [15, 16, 20, 21]. Few of them did so by establishing a metric called Codon stabilization coefficient(*CSC*), which is a pearson correlation coefficient between the occurrence of each codon in the transcript with the transcript’s half-life. The optimal codons were found to have positive correlation with the mRNA half-life while non-optimal codons were rendered negative [15, 16]. Analysis using human and mammalian cell lines showed similar results, where they synthesized synthetic constructs enriched in both optimal and non-optimal codons to study their half-lives in real-time [20, 21]. But an mRNA transcript is an ensemble of all the 61 codons, where the combination of *NOC* and *OC* can be differentially distributed along the length of the transcripts exerting differential effects on its stability.

Due to above shortcomings, we introduced *CI*, which is a normalized difference between the non-optimal and optimal codon content of a transcript with its value ranging from −1 to +1. We have classified *CI* range into three different groups, (i) higher amount of optimal codons within a transcript, where the *CI <* 0.1, (ii) higher amount of non-optimal codons within a transcript, where *CI >* 0.1, and (iii) an intermediate between optimal and non-optimal codons, where 0.1 *CI* 0.1. In accordance with previous results [21, 20, 16, 15], we found that the stability of the transcripts increases with increase in amount of optimal codon (Fig. 1A). Transcripts with higher abundance of optimal (*CI <* − 0.1) and non-optimal codons (*CI >* 0.1), show the highest and the lowest median half-lives 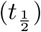 respectively (Fig. 1A). The distributions differ significantly when compared with multi-sample Kruskal Wallis (*KW*) test for median (*KW p* = 9.16E-185)(Fig. 1A). It should be noted that the codons are not always equally distributed along the length of the transcript as there can be long stretches of optimal or non-optimal codons. We embark on introducing both frequency (*f*) and length of *NOC* and investigate their effects on mRNA stability both separately and combined in the following sections.

**Figure 1:**
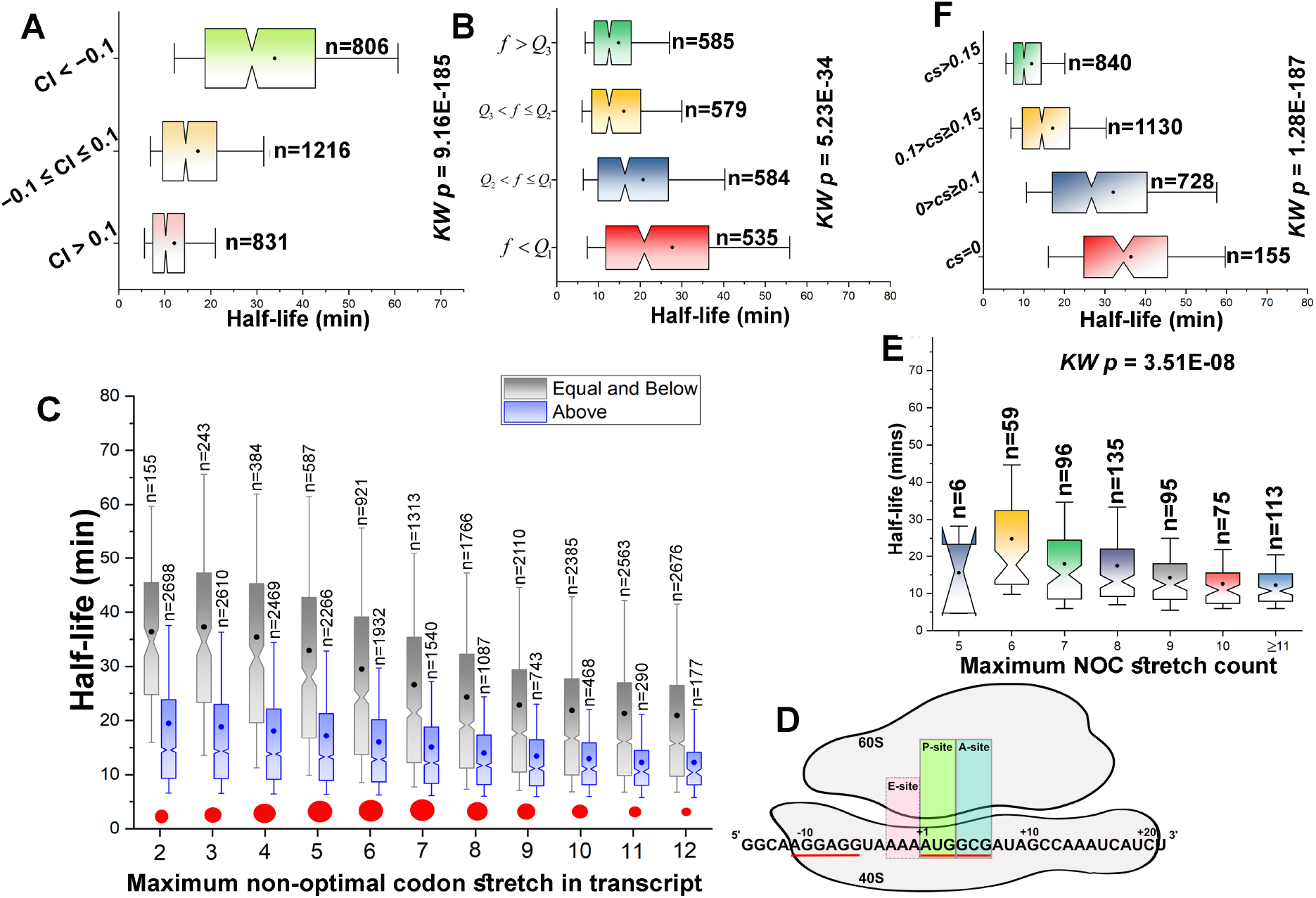
A) Effects of codon content index (*CI*) on transcript half-life. Transcripts with higher value of *CI*, indicating higher amount of non-optimal codon show reduced half-life. (B) Notched box plot showing the effect of frequency(*f*) of *NOC* stretch *≥* 3 on the transcript half-life. Transcript with higher frequency of *NOC* stretch show reduced mRNA half-life. *f* is classified based on Quartile (*Q*) (C) A comparative notched box plot between transcripts having non-optimal codon stretch equal and below (grey) and above (blue) a threshold. The transcripts above the threshold (blue) always show a stunted half-life. Mann-Whitney (*MW*) test is performed between the two groups, and the *p-value* is denoted by red circle. We observed maximum half-life difference between groups have a continuous stretch of 5-7 non-optimal codons. (D) A model diagram representing the span of mRNA along the ribosomal tunnel, which can fit around 31 nucleotides [35]. Post the A-site, the tunnel spans through a region of 15 nucleotides (5 codons).(E) Transcript within a fixed frequency range (*Q3 < f < Q2*) are classified based on their maximum *NOC* stretch count and their half-lives are plotted. Half-life decreases gradually with increase in *NOC* stretch (F) A scoring metric (Cumulative score, *cs*) which takes account of *f* and length of the *NOC* within a transcript. Longer stretch *NOC* are given a higher weightage in contrast to lesser stretch during scoring. Transcript’s half-life gradually decreases with increase in median *cs*. Multi-sample Kruskal Wallis (*KW*) test is performed and the *p-value* is provided to test whether the distributions differ significantly. The plots are generated using experimental half-life data by Miller and his group [23]. Similar calculations for the other two half-life datasets [15, 24] are performed and their respective *p-values* are provided in Data S3.

#### 3.1.1 Higher the frequency of Non-optimal codon stretches, lower the transcript stability

We have estimated the frequency (*f*) of *NOC* stretch length(consecutive presence of *≥* 3 *NOC* ‘s) along the transcript irrespective of the stretch length. The frequency(*f*) of *NOC* stretch ranges from 1 to 178 with a mean 27.6 across 2283 transcripts (Fig. S1A, Data S2). With such a wide range of *f*, we classified *f* into four sets (*G1,G2,G3 and G4*) based on their quartile (*Q*). The sets are (i) *f <Q1* (*G1*), (ii)*Q1≤ f <Q2* (*G2*), (iii) *Q2≤ f <Q3* (*G3*) and (iv) *f≥ Q3* (*G4*). Half-life comparisons between these four sets showed decreased transcript stability with an increase in frequency of *NOC* stretches (Fig. 1B). Analysis revealed medians of 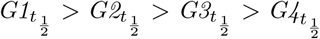, and their distributions show statistical difference (*KW p =*5.23*E −* 34).

In another analysis, half-lives of transcripts are compared belonging to (i) *G1* and *G2,G3 and G4*, (ii) *G1,G2* and *G3 and G4*, (iii) *G1,G2 and G3* and *G4*. The results reveal that transcripts belonging to lower quartile groups have a higher median half-life compared to transcripts belonging to higher quartile groups (Fig. S1B). As a result, we can see loss in mRNA stability with an increase in frequency of *NOC* stretches.

Codon optimality is the result of differential *aa*_*tRNA*_ concentrations within the cellular pool, which result in a non-uniform decoding rate for different transcripts [11, 30]. Optimal codons are rapidly incorporated due to the higher concentration of *aa*_*tRNA*_ in the cellular pool, and over-represented in highly expressed genes. In a study, translation speed were compared between an optimized and non-optimized coded mRNA of same length (1.6kb). It is observed that codon optimized mRNA is translated 1.5 times faster than that its non-optimized control [31]. Real time monitoring of translation by a fluorescence-based method revealed a speed of 4.9 codons per second compared to 3.1 codons per second for a non-optimized one [32]. Frequency of *NOC* stretches shows a high positive correlation (*Pearson r* =0.95, *p*=0) translation time (seconds) of yeast transcriptome [33] (Fig. S1C). Translation time of transcripts having *NOC* stretches at the beginning have higher median translation times compared to transcripts with no *NOC* stretches (Fig. S1D).

Therefore, in a way a non-optimal codon does dictate the ribosomal speed over the transcript due to its limited concentration of 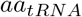 within the cellular pool. A higher frequency of non-optimal codon stretch in a transcript often restricts ribosomal movement leading to a very low translation rate with intermittent stoppages. A restricted or no ribosomal movement along the transcripts in any of the stretches, triggers Ccr4-Not complex to disengage transcripts and pass it to the cellular degradation machinery [10].

#### 3.1.2 Longer non-optimal codon stretches facilitate mRNA decay

We have calculated the *NOC* stretches for whole yeast transcriptome and mapped them along with their half-life 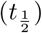 values. Our analysis includes only those transcripts, for which the maximum *NOC ≥*3. Transcript stability gradually decreases with an increase in maximum *NOC* stretches. The half-life distributions were found to be significantly different (*KW p* = 8.41E-72)(Fig. S1E). Longer *NOC* stretch length neccesitates for more time owing to the reduced concentration of *aa*_*tRNA*_ within the cellular pool.

The translational speed is limited to a bare minimum, or will lengthen the ribosomal halt period. This is an opportune moment for Ccr4-Not complex to disengage transcripts and pass them to the cellular degradation machinery.

Our next aim is to track down the exact *NOC* stretch length for which the transcript half-life will differ maximally. In order to examine this, we classified the transcripts into two groups, (i) Equal and below, and (ii) Above a *NOC* stretch length and checked the difference in half-life between the two groups. Each of the pairs is tested statistically using pairwise Mann-Whitney U test (*MW*) for medians. The *NOC* stretch length is varied from 3 to 12. In accordance with previous analysis, we found transcript belonging to the group above are less stable than the transcripts belonging to the equal and below for that same pair. The *MW* p-value difference between the pairs gradually increases from 2, peaks at *NOC* stretch length of 5,6, and 7, and again steadily decreases further above (Fig. 1C).

About 31 nucleotides can be accommodated in the groove between 40S and 60S subunit of ribosome for translation [34, 35]. The 40S subunit recognizes the Shine Dalgarno/Kozak sequence (−10 upstream of mRNA) and places itself in such a way that the P-site is stationed on the +1(AUG) start codon. The A-site now places itself from +4 to +6 position of the transcript’s coding region. Post A-site, the mRNA spans for another 15 nucleotides (+21 of the transcript’s coding region) into the ribosomal translation tunnel (Fig. 1D). Our result reveals maximum half-life difference with a peak of *MW-p*= 3.84*E −* 81 and 2.31*E −* 81 for a *NOC* stretch length of 5 and 6 respectively, which is about 15-18 nucleotides. Beyond P-site, the mRNA spans from +4 to +21 (coding region) into the ribosome, which is around 18 nucleotides (6 codons) region and beyond A-site, the mRNA spans for 15 nucleotides (from +6 to +21 of the coding regions, 5 codons) region. So, a *NOC* stretch of 15-18 nucleotides, will drastically slower the incorporation speed into the A-site and since, eviction of [ieq] from E-site does not depend on induction of *aa*_*tRNA*_ onto A-site, so a prolonged exposure of empty E-site and A-site is an indication of slow decoding kinetics of ribosomes which can be sensed by Ccr4-Not complex to trigger mRNA release and degradation [10].

#### 3.1.3 Higher frequency and longer stretch length of *NOC* cumulatively degrade mRNA faster

So far, we have observed *f* and *NOC* adversely affect mRNA stability. Next, we embark on checking their cumulative effect on mRNA half-life. For doing so, we regenerate Fig. S1B for (*G1*), (*G2*), (*G3*) and (*G4*) individually. So, for each frequency group, we classify the transcripts based on their maximum *NOC* stretch length. The median half-life decreases steadily with increase in maximum *NOC* stretch length for *G1,G2,G3* and *G4* (Fig. 1E, Fig. S1F). The distributions are tested using multi-sample *KW* test (*p-value*= 3.51*E−* 08) (Fig. 1E).

Furthermore, we divide the transcript into two groups, (i) *f <Q1* and *NOC* stretch length of 3 (lower extreme) and (ii) *f >Q3* and *NOC* stretch length *≥* 12 (higher extreme). Transcript belonging to the lower extreme group have a median of 37.46 minutes compared to 11.65 minutes for transcripts belonging to the higher extreme group. Both these distributions are tested using pairwise *MW* -test with a *p-value* of 3.70*E −* 19 (Fig. S1G).

Parallely, we mapped transcripts belonging to all four frequency groups, to their *NOC* stretch length distribution. Amount of long length *NOC* stretches (*≥* 5) have increased gradually within transcripts belonging to higher frequency groups from low frequency groups, while there is a very negligible difference between shorter *NOC* stretches (*≤* 4)(Table S1). Hence, longer length stretches are characteristic of transcripts belonging to higher frequency groups, hence contributing to their reduced stability and half-life.

#### 3.1.4 A combined metric for evaluation of both *NOC* stretch length and frequency for a transcript

Since both *f* and length of *NOC* stretch proved to be important for regulating mRNA half-life, we have introduced cumulative score (*cs*), which considers both the parameters for a single transcript. The metric *cs* includes a weightage factor (*k*) which is varied from 0.1 to 10. Since the power of *f* is kept constant, so lowering *k* below 1, increases the contribution of *f* on the *cs* and vice versa. We mapped *cs*-score of transcripts against their half-life. Transcripts belonging to higher *cs*-score group, tends to be less stable and have a lower median half-life compared to transcripts belonging to lower *cs*-group for all values of *k* (Fig. 1E for *k* =2, Fig. S1H for rest value of *k*, Data S2). The distributions are statistically compared using multi-sample Kruskal-Wallis and the *p*-values are given in Table S2. We can see the statistical difference gradually increases with *k* from 0.1, until it becomes almost static beyond *k* of 0.33. Hence, we can say that both frequency (*f*) and *NOC* stretch length share their effects towards mRNA stability. However, we cannot conclude whether *f* or *NOC* has any comparatively higher effect than the other.

### 3.2 Combinatorial role of IUS and Codon composition on mRNA stability

The primary function of mRNA is to translate the DNA information to amino acids of the protein via translation. mRNA are averted to degradation once the fidelity or efficiency of translation is compromised. Elimination of mRNA from the translational pool can occur through interaction with the Ccr4-Not complex [10] or Dhh1 [36] by the factors mentioned above. The transcripts released for degradation are first broken down into smaller entities by endonucleases, which are then scavenged by exonucleases [37]. Here, *IUS* can play an integral role providing amenable platform for endonuclease engagements. So, we integrate *IUS* data with *cs*, to find out their combinatorial role on the mRNA half-life.

### 3.2.1 *IUS* in combination with higher *cs* score facilitate easy decay of transcript

*IUS* destablizes mRNA by providing suitable attachment sites for endonuclease action [1]. Transcripts released from the translational system owing to prolonged stopagges are scavenged by these endonucleases for degradation. Therefore, we hypothesize that both of these factors (*cs* and *NOC*) will act as agonists in lowering the half-life and stability of mRNA.

In order to investigate the said hypothesis, we have classified transcripts based on their *cs* score based on the median(*M*_*cs*_) into two groups, (i) *cs< M*_*cs*_ (Fig. 2A) and (ii) *cs ≥ M*_*cs*_ (Fig. 2B). Furthermore, each of these groups (having same range of *cs)* are classified on the basis of *IUS* (*ξ*). The value of *ξ* is classified according to the quartile(*Q*) into three separate groups, (i) *ξ < Q*_1_, (ii) *Q*_1_ *≥ ξ ≤ Q*_3_ and (iii) *ξ > Q*_3_. As said, transcripts with higher amount *IUS* (*ξ > Q*_3_) showed reduced stability, and transcript with *ξ < Q*_1_ showed maximum stability for each *cs* group (Fig. 2A,B). Distributions are tested using *KW* multi-sample test for median and are found to be significant. The plot for the structured threshold of 60% is shown in Figs. 2A and B, while the plots using all the other structured thresholds (30%, 40%, 70% nucleotides having PARS *>* 0) are given in Fig. S2A.

**Figure 2:**
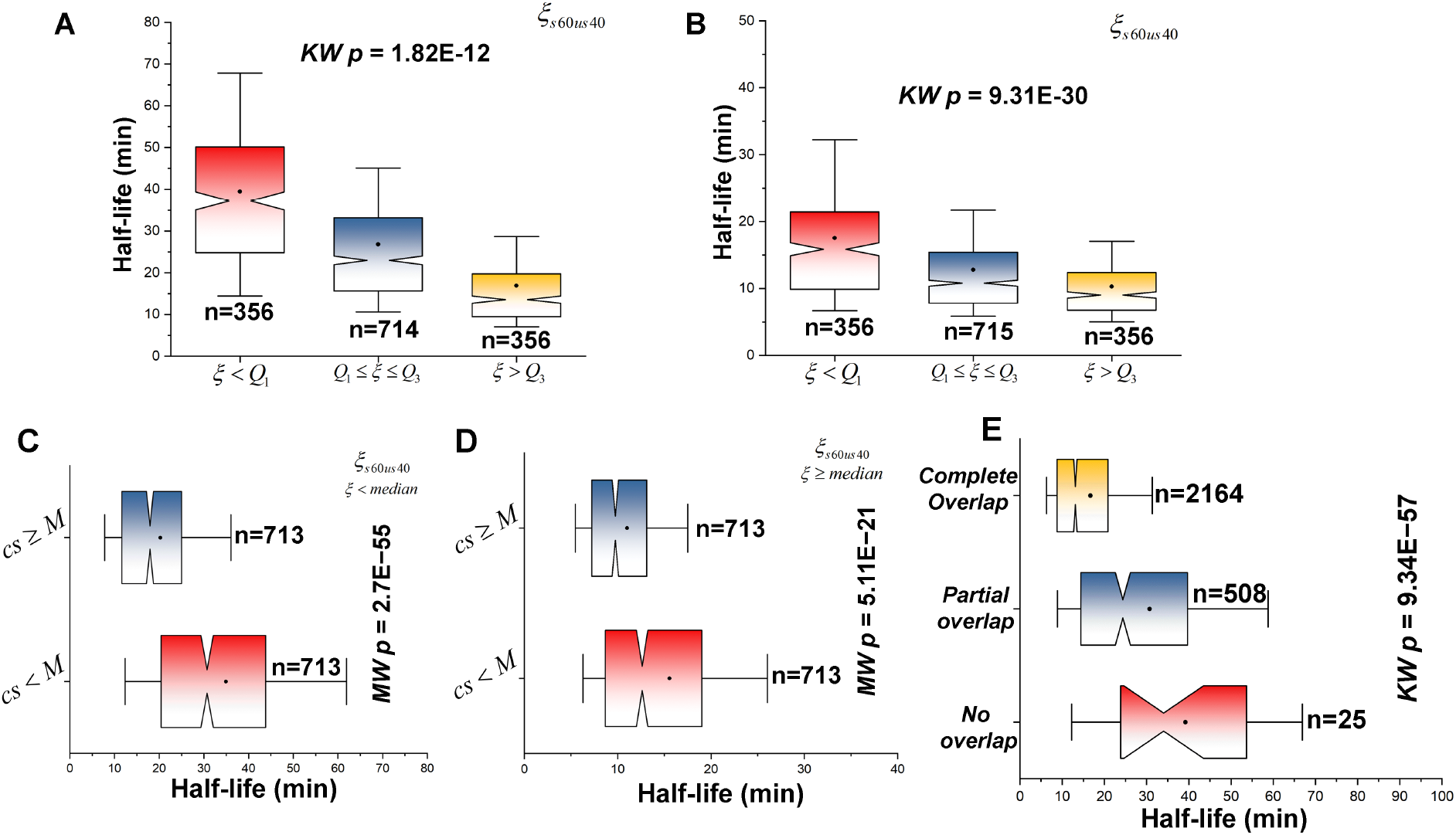
Combinatorial effect of *IUS* (*ξ*) and *cs* on transcript half-life. A) Transcript with a *cs <* Median(*M*) are classified based on their *IUS* (*ξ*). (B) Transcript with a *cs ≥* Median(*M*) are classified based on their *IUS* (*ξ*). For both (B) and (C), *ξ* is classified based on quartile (*Q1* and *Q3*). (C) For a fixed range of *ξ* (*ξ < M*), transcripts are classified based on their *cs*. (D) For a fixed range of *ξ* (*ξ ≥ M*), transcripts are classified based on their *cs. cs* is classified based on the median value (*M*). Transcripts with a higher value of *cs*, show reduced half-life within a fixed *ξ* range. (E) Half-life of transcripts are compared having complete, partial and No overlap between *IUS* and *NOC*. For all plots involving *ξ*, plots are only generated for *ξ*_*s*60*us*40_. For (A), (B) and (E), multi-sample Kruskal Wallis (*KW*) test is performed and the *p-value* is provided to test whether the distributions differ significantly. For (C) and (D), two-sample Mann-Whitney test is performed and the *p-value* is provided to test the same. The plots are generated using experimental half-life data by Miller and his group [23] using *ξ*_*s*60*us*40_. The plots for other structured-unstructured threshold of *IUS* is present in Fig S2. Similar calculations for the other two half-life datasets [15, 24] are performed and their respective *p-values* are provided in Data S3.

A reverse analysis is also performed by classifying the *ξ* values of the transcripts into two groups, which are further classified into two groups, each based on their *cs* score. The transcripts are initially divided as (i) *ξ < M*_*ξ*_ (Fig. 2C) and (ii) *ξ ≥ M*_*ξ*_ (Fig. 2D). In addition, each of the mentioned groups is subdivided according to their *cs* as (i) *cs< M*_*cs*_ and (ii) *cs≥ M*_*cs*_. No surprise, we find that the transcript belongs to the group *cs< M*_*cs*_ shows highest median half-life compared to *cs ≥ M*_*cs*_ which show the least median half-life for both the classifying *cs* group (Fig. 2). The half-life distributions of the two *cs* groups are found significant when tested using pairwise *MW* -test for median (Figs. 2C,D, S2B). Similar to the last analysis, the *ξ* classification for structured threshold 60% is shown in Fig. 2C and D, while for the rest(30%, 40%,70% nucleotides having PARS *>* 0) is shown in Fig. S2B.

The above result is an indicatation of the importance of *IUS* within the released transcripts for degradation. Transcripts released from translation system for degradation can be left in the cytosol or sequestered into P-bodies. Meanwhile, both places contain an ensemble of decapping and endonucleolytic cleavage factors [38, 39, 40]. Endonucleases will break down larger transcripts into smaller entities, while 5’-decapping factors can remove 5’-cap from mature mRNA so that it can be easily abducted by various exonucleases for complete degradation. In case of endonuclease cleavage, presence of *IUS* can prove amenable for enzyme-substrate engagement platform. The frequency of *IUS* will increase the probability of these engagements breaking down the large transcripts into smaller entities for exonuclease capture.

#### 3.2.2 Transcripts with overlapped *NOC* stretches and *IUS* regions degrades faster

Next, we want to find the precise location of *NOC* stretches and *IUS* segments and their distribution. We have classified their position along their transcript as (i) no overlap, (ii) partial overlap where both either 3’ end or 5’ end overlaps with each other and (iii) complete overlap, where one stretch is positioned within another stretch (Data S1 and S2). We found both *NOC* stretches and *IUS* region mutually lowers the transcript’s half-life. Complete overlap between *NOC* stretches and *IUS* facilitates easy release from the translational machinery and availability of cellular endonuclease speed up the capturing probability via the *IUS* segments. Hence, they show least transcript stability and median half-life (Fig. 2E, Fig. S2C). The transcripts containing no overlap among *NOC* and *IUS* show most stability (Fig. 2E, Fig S2C). The three groups are tested statistically using multi-sample *KW* test for median (*KW* p = 9.34E-57). Therefore, these overlapping regions act as premiere hotspots which will smoothen the passageway for the mRNA degradation process.

### 3.3 Ribosome association offers stability to *NOC* and *IUS* containing tran-script

Translational rate depends on the ribosomal occupancy during the elongation phase of translation along the length of the mRNA. Ribosomes are placed at regular intervals to form a polysome, thus governing the rate of protein synthesis [41]. In addition to regulating the rate of protein synthesis, ribosomal association also stabilizes the mRNA transcript by a significant level by protecting mRNA molecules from engagement of the exosomal complex [42]. Nearly 25% increase in transcript stability is observed with around 78% increase in ribosomal density [22]. There are several studies which indicates NMD targeted transcripts are more vulnerable to degradation with their monosomal form compared to their polysomal forms [43, 44, 45]. Additionally, we have shown that mRNA sequestration with RNA Binding proteins (*RBP*) imparts stability to the transcript [1]. But a genome scale relationship between ribosome, which is a macromolecular protein complex and mRNA stability remains unclear till date.

To investigate this, we used experimental data from ribosomal profiling (*rf*) from three different datasets (which differ in their extraction methods of total RNASeq count) [27] (Data S1). The *rf* value indicates the ribosomal density per kilobases of the transcript. So, a higher *rf* value of a transcript signifies higher concentration of ribosomes residing on that transcript. We plotted the half-life value of the transcripts against their respective *rf* s. Our result reveals that the increased ribosomal density aids in increased transcript stability (Fig. 3A, Fig. S3A). The half-life distributions are found statistically significant using multi-sample Kruskal-Wallis test (*KW p-value* = 0)(Fig. 3A, Fig. S3A).

**Figure 3:**
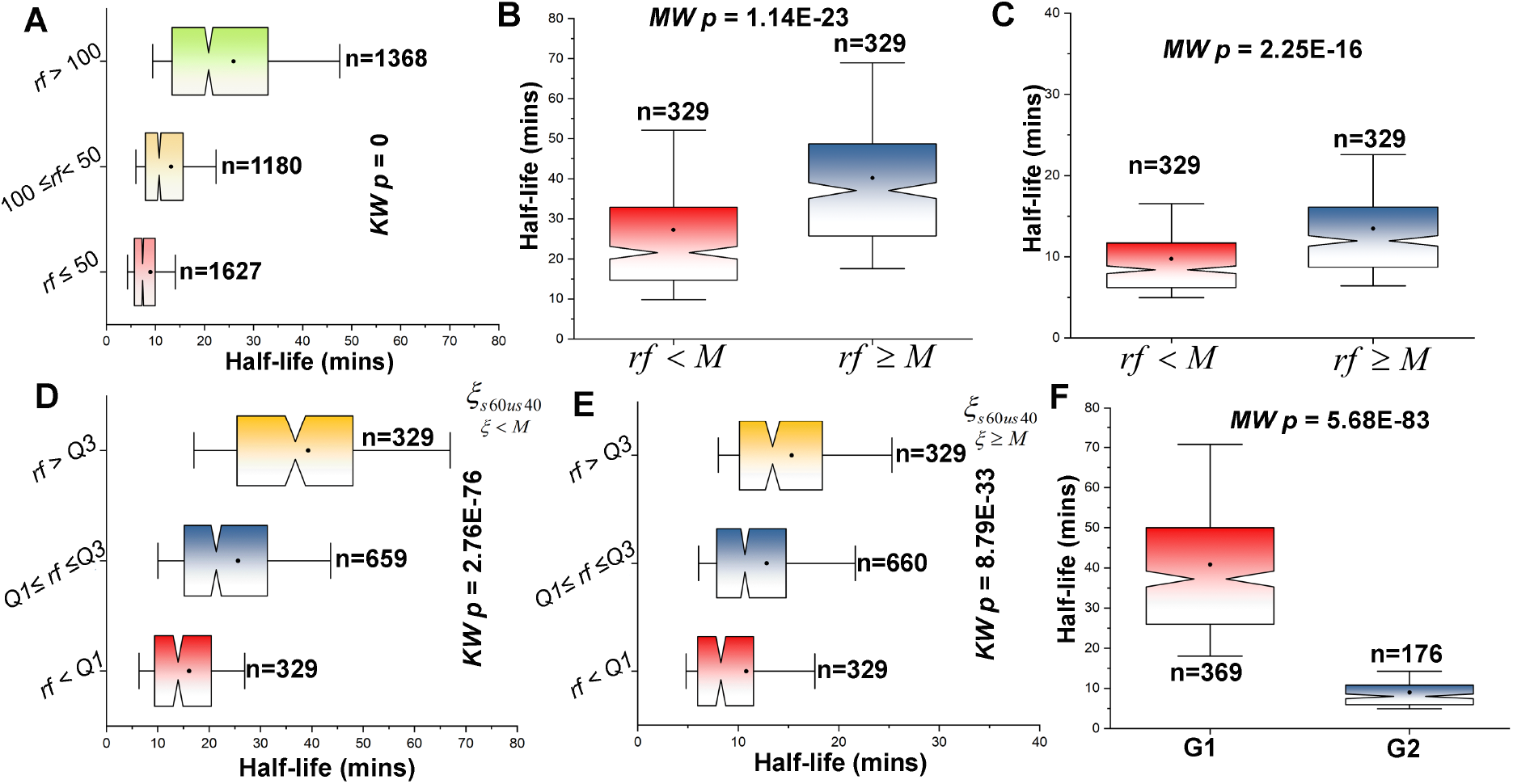
Effects of Ribosomal association on transcript’s half-life (A) Transcripts are classified based on their *rf* value, and their half-life are compared. Transcripts with higher rf value offers more stability to the transcript and hence enhance its half-life. (B) and (C) For a fixed *cs*, trancript’s having (C) *cs <* Median(*M*) and (D) *cs ≥* Median(*M*) are classified based on their *rf* value. Among them transcripts with *rf >* Median(*M*) show higher stability for each of the representative group. (D) & (E) For a fixed *ξ* (*ξ < M*) (C) and (*ξ ≥ M*) (D), transcripts are classified based on their *rf* value. Transcript’s stability gradually increases with increase in *rf* value. (F) Transcripts are classified based on two extreme conditions, (i) *cs < Q1, ξ < Q1* and *rf > Q3* referred as G1 and (ii) *cs > Q3, ξ > Q3* and *rf < Q1* referred as G2. The half-life between these two extremes is compared. For (A), (D) and (E), multi-sample Kruskal Wallis (*KW*) test is performed and the *p-value* is provided to test whether the distributions differ significantly. For (B), (C) and (F), two-sample Mann-Whitney test is performed and the *p-value* is provided to test the same. For *rf* data, plots are generated only for Ribominus datasets. For *ξ*, plots are only generated for *ξ*_*s*60*us*40_. The plots are generated using experimental half-life data by Miller and his group [23] using only Ribominus footprinting data. The plot for rest of *ξ* structured-unstructured threshold for the rest of footprinting data (Unselected and Dynabeads) is shown in Fig S3. Similar calculations for the other two half-life datasets [15, 24] are performed and their respective *p-values* are provided in Data S3.

We have already established that *cs* is a metric of quantifying both *NOC* stretch length and stretch count of a transcript, and a higher *cs* score renders less stability. Similarly, a higher ribosomal density of the transcripts renders more stability to the transcript and increased half-life. Therefore, we look to overlay these two parameters in order to understand the effect of each other on the transcript.

To do so, we have mapped *cs, rf* and half-life values of the transcript. The transcripts are classified as the least stable (*cs > Q*_3_) and most stable (*cs < Q*_1_). The transcripts falling under the least stable category are further classified based on their median *rf* value. The two distributions show significant difference (*MW p-value* = 1.14*E−* 23) (Fig. 3B), with the group equal and above median *rf* shows higher median half-life (38.08 min) compared to the group below median *rf* showing a median half-life of 21.56 min. An identical calculation is also performed for the most stable transcripts, and a similar result is found (*MW p-value* = 2.25*E −* 16)(Fig. 3C, Fig. S3B). The median half-life for the group equal and above median *rf* and below median *rf* is 11.95 and 8.4 min, respectively. The median half-life values and the statistical significance between the most and least stable groups indicate the magnitude of instability exerted by the presence of non-optimal codon.

Ribosomes are a macromolecular complex which can greatly protect unstructured regions of the transcript from the various endonucleases in the cellular pool. In our previous work, we found that transcripts with higher fractions of *IUS* buried into RBP sites have higher stability compared to lower fraction transcripts [1]. In a similar manner, we enquired whether ribosomal association can enhance stability of transcripts with *IUS*. So, we classified *ξ*_*s*60*us*40_ *≥* based on the median (M) of their distribution into two groups. They are (i) *ξ*_*s*60*us*40_ *< M* and (ii) *ξ*_*s*60*us*40_ ≥ *M*. These two groups are further classified based on their *rf* value into three groups based on the quartile (*Q*) value of the distribution. Our analysis reveals transcripts with highest ribosomal density value (*rf ≥ Q*_3_), show the highest stability compared to the lower groups. The difference in half-life is tested using multi-sample Kruskal-Wallis and is statistically significant in both the cases (Fig. 3D,E, Fig S3C-F). For the rest of the structured threshold (30%, 40%,60%,70% nucleotides having PARS *>* 0) and extraction methods, the plots are presented in Fig. S3C-F.

### 3.4 Comparison between two transcript groups having extreme parameter values

We have illustrated cumulative score(*cs*), Internal unstructured region (*IUS*) and ribosomal footprinting (*rf*) both in their singular and combinatorial form can act as regulators of mRNA stability. Lastly, we accumulate all the three parameters and check how the half-life differs between the two extremes. Here, we define the two extremes as (i) transcripts having all the parameters value which will render them stabilising (*G1*) and (ii) transcripts having all the parameter values which will render them destabilising (*G2*). *G1* include transcripts with (i)*cs < Q1*, (ii) *ξ < Q1* and (iii) *rf > Q3* while *G2* include transcripts with (i) *cs> Q3*, (ii) *ξ >* Q3 and (iii) *rf < Q1. G1* has a median half-life of 37.25 min compared to *G2* which has a median half-life of 8.035 min. Both the distributions are significantly different when tested with pairwise Mann-Whitney (*MW-p* value = 5.68*E −* 83)(Fig. 3F, S3G-H).

### 3.5 Sequence and structural features, along with ribosomal binding in mRNA regulate its co-translational degradation

Since mRNA is an ensemble of *NOC* and *OC* translated by multiple ribosomes, different regions of the transcripts experience different translation rates. Optimal codons can increase translational efficiency with their large reservoir of cognate tRNAs. Non-optimal codons often slow down translational speed (Fig. S1D,E), leading to ribosomal collisions, frequent halts and ultimately release and degradation of transcripts when it is beyond recovery [11]. Moreover, we found correlation between translation rate (Length of transcript/Translation time) with *CI* (*Pearson r* =-0.92, *p*=0) (Fig. S4A) and *cs*(*Pearson r* =-0.82, *p*=0)(Fig. S4B). These strong negative correlations suggest that transcripts with higher amount of non-optimal codons experience slow ribosomal movement. Slow translational speed is a result of slow peptidyl transfer, prolonged vacancy of A-site of ribosomes which can be sensed by Ccr4-Not or Dhh1. These protein complexes are associated with numerous deadenylation and decapping holoenzymes, which will define direct pathway of these transcripts to the degradation machineries giving them a low characteristic half-life. Thus codon optimality influences the translation rate and half-life of mRNA which in turn regulates the gene expression and protein levels in cells [12]. However, translation rate poorly correlates with mRNA half-life(*Pearson r* = 0.47, *p*=2.72E-146). A weak correlation indicates that both translation rate and mRNA half-life are not primarily dictated by each other. Till now, we have explored few translation-linked regulatory factors (described in this study) which also governs mRNA half-life. But in reality, there are a myriad of factors which regulate translation rate and half-life in different ways. One such factor is mRNA secondary structure. Ribosomal progression is often hampered by local secondary structures within mRNA which in turn can hamper the translation rate. However, local secondary structures rarely affect degradation since almost all degradation machineries are equipped with powerful motor proteins that can unwind and channel the transcript for decay [46]. So, the fuzzy positive correlation between translation rate and mRNA half-life can be account of many more such instances.

Releasing the transcripts from the translational system owing to their reduced rate always does not ensure their degradation. Ccr4-Not complex is associated with 5’-decapping and 3’-poly-deadenylation enzymes, which can make these transcripts defenseless towards any degradation machinery [7, 8, 9]. It is to note that transcripts destined for decay are first broken down into smaller entities by cellular endocnucleases present in the cytosol, P-bodies or various stress granules and later on these are scavenged by directional exonucleases [37]. So, after the release of transcripts, endonuclease action prefers presence of internal unstructured segments(*IUS*) for enzyme-substrate engagements. To understand whether the release and capture mechanism works in an additive way, we combined *cs*-score, which is a unified metric for both frequency and stretch length of *NOC*, with *IUS*. We found decreased stability for transcripts with higher amount of *IUS* and *NOC* stretches (Fig. 2, Fig. S2).

We have previously reported that association of mRNA with RNA-binding protein (*RBP*) protects the mRNA from being captured by degradation machineries by concealing the enzyme attachment regions [1]. Here, we have used ribosomal footprinting data to understand whether polysome formation can stabilise a transcript even in the presence of *NOC* stretches and *IUS* within transcripts. Ribosome, being a large macromolecular complex can bury large parts of the transcripts which are susceptible to nuclease action and can elevate their half-lives (Fig. 3, Fig. S3).

In addition to our previous work, this analysis aims to uncover more factors that could regulate mRNA stability during translation. In future, these parameters can prove valuable in application-oriented research for designing synthetic transcripts with maximized efficacy for mRNA based medication, drug delivery systems, and vaccines [47].

## 4 Conclusion

In this study, we embark on understanding the reason for mRNA release and degradation during translation by using multi-strata data (sequence, structural, and ribosomal binding data) from different sources. We have used relative abundance of *NOC* within a transcript, *NOC* stretch frequency and length, *IUS* and ribosomal footprinting to understand their effects on mRNA stability.

i. Higher fraction of non-optimal codons(*NOC*) within transcripts (*CI*) leads to reduced stability and half-life.
ii. We observed, (a) the presence of longer *NOC* stretch length, (b) higher frequency of *NOC* stretches facilitates release of transcripts from the translational machinery to the degradation machinery and has reduced stability.
iii. *IUS* within transcript makes the transcript susceptible to endonuclease attachments and degradation after being released from the translation system due to *NOC*, and hence reducing it shelf life.
iv. Ribosomal attachment has been shown to stabilize the transcript even in the presence of destabilizing factors like *NOC* stretch and *IUS*.

## Author Contribution

S.B. and S.K. designed the research; S.B. implemented the computational methodology, S.B., R.M. and

S.H. performed research; all analysed data and wrote the paper.

## Acknowledgement

S.B is supported by GenML Solutions (OPC) Pvt. Limited. S.H is supported by DBT-BINC SRF fellowship (Fellow Number: DBT-BINC/2017/CU/12). We would also like to acknowledge Dr. Runa Sur and Dr. Jayanta Mukhopadhyay for their valuable comments on the manuscript.

## References

[1] Sudipto Basu et al. “Genome-scale molecular principles of mRNA half-life regulation in yeast”. In: The FEBS Journal 288.11 (2021), pp. 3428–3447.

[2] Wenqian Hu et al. “Co-translational mRNA decay in Saccharomyces cerevisiae”. In: Nature 461.7261 (2009), pp. 225–229.

[3] Saverio Brogna and Jikai Wen. “Nonsense-mediated mRNA decay (NMD) mechanisms”. In: Nature structural & molecular biology 16.2 (2009), pp. 107–113.

[4] A Alejandra Klauer and Ambro van Hoof. “Degradation of mRNAs that lack a stop codon: a decade of nonstop progress”. In: Wiley Interdisciplinary Reviews: RNA 3.5 (2012), pp. 649–660.

[5] Yuriko Harigaya and Roy Parker. “No-go decay: a quality control mechanism for RNA in translation”. In: Wiley Interdisciplinary Reviews: RNA 1.1 (2010), pp. 132–141.

[6] Martine A Collart. “The Ccr4-Not complex is a key regulator of eukaryotic gene expression”. In: Wiley Interdisciplinary Reviews: RNA 7.4 (2016), pp. 438–454.

[7] Ingmar B Schäafer et al. “Molecular basis for poly (A) RNP architecture and recognition by the Pan2-Pan3 deadenylase”. In: Cell 177.6 (2019), pp. 1619–1631.

[8] Eugene Valkov et al. “Structure of the Dcp2–Dcp1 mRNA-decapping complex in the activated conformation”. In: Nature structural & molecular biology 23.6 (2016), pp. 574–579.

[9] Roy Parker. “RNA degradation in Saccharomyces cerevisae”. In: Genetics 191.3 (2012), pp. 671–702.

[10] Robert Buschauer et al. “The Ccr4-Not complex monitors the translating ribosome for codon optimality”. In: Science 368.6488 (2020).

[11] Gavin Hanson and Jeff Coller. “Codon optimality, bias and usage in translation and mRNA decay”. In: Nature reviews Molecular cell biology 19.1 (2018), pp. 20–30.

[12] Jan-Hendrik Troösemeier et al. “Optimizing the dynamics of protein expression”. In: Scientific reports 9.1 (2019), pp. 1–15.

[13] Laura Jeacock, Joana Faria, and David Horn. “Codon usage bias controls mRNA and protein abundance in trypanosomatids”. In: Elife 7 (2018), e32496.

[14] Tamir Tuller et al. “Translation efficiency is determined by both codon bias and folding energy”. In: Proceedings of the national academy of sciences 107.8 (2010), pp. 3645–3650.

[15] Vladimir Presnyak et al. “Codon optimality is a major determinant of mRNA stability”. In: Cell 160.6 (2015), pp. 1111–1124.

[16] Yuriko Harigaya and Roy Parker. “Analysis of the association between codon optimality and mRNA stability in Schizosaccharomyces pombe”. In: BMC genomics 17.1 (2016), pp. 1–16.

[17] Gréegory Boëel et al. “Codon influence on protein expression in E. coli correlates with mRNA levels”. In: Nature 529.7586 (2016), pp. 358–363.

[18] Ariel A Bazzini et al. “Codon identity regulates mRNA stability and translation efficiency during the maternal-to-zygotic transition”. In: The EMBO journal 35.19 (2016), pp. 2087–2103.

[19] Yuichiro Mishima and Yukihide Tomari. “Codon usage and 3’ UTR length determine maternal mRNA stability in zebrafish”. In: Molecular cell 61.6 (2016), pp. 874–885.

[20] Qiushuang Wu et al. “Translation affects mRNA stability in a codon-dependent manner in human cells”. In: elife 8 (2019), e45396.

[21] Megan E Forrest et al. “Codon and amino acid content are associated with mRNA stability in mammalian cells”. In: PloS one 15.2 (2020), e0228730.

[22] Shlomit Edri and Tamir Tuller. “Quantifying the effect of ribosomal density on mRNA stability”. In: PLoS One 9.7 (2014), e102308.

[23] Christian Miller et al. “Dynamic transcriptome analysis measures rates of mRNA synthesis and decay in yeast”. In: Molecular systems biology 7.1 (2011), p. 458.

[24] Benjamin Neymotin, Rodoniki Athanasiadou, and David Gresham. “Determination of in vivo RNA kinetics using RATE-seq”. In: Rna 20.10 (2014), pp. 1645–1652.

[25] J Michael Cherry et al. “SGD: Saccharomyces genome database”. In: Nucleic acids research 26.1 (1998), pp. 73–79.

[26] Nicholas T Ingolia. “Ribosome footprint profiling of translation throughout the genome”. In: Cell 165.1 (2016), pp. 22–33.

[27] David E Weinberg et al. “Improved ribosome-footprint and mRNA measurements provide insights into dynamics and regulation of yeast translation”. In: Cell reports 14.7 (2016), pp. 1787–1799.

[28] Michael Kertesz et al. “Genome-wide measurement of RNA secondary structure in yeast”. In: Nature 467.7311 (2010), pp. 103–107.

[29] øyvind Hammer, David AT Harper, Paul D Ryan, et al. “PAST: Paleontological statistics software package for education and data analysis”. In: Palaeontologia electronica 4.1 (2001), p. 9.

[30] Mridusmita Saikia et al. “Codon optimality controls differential mRNA translation during amino acid starvation”. In: Rna 22.11 (2016), pp. 1719–1727.

[31] Chien-Hung Yu et al. “Codon usage influences the local rate of translation elongation to regulate co-translational protein folding”. In: Molecular cell 59.5 (2015), pp. 744–754.

[32] Xiaowei Yan et al. “Dynamics of translation of single mRNA molecules in vivo”. In: Cell 165.4 (2016), pp. 976–989.

[33] Marlena Siwiak and Piotr Zielenkiewicz. “A comprehensive, quantitative, and genome-wide model of translation”. In: PLoS computational biology 6.7 (2010), e1000865.

[34] Joan Argetsinger Steitz. “Polypeptide chain initiation: nucleotide sequences of the three ribosomal binding sites in bacteriophage R17 RNA”. In: Nature 224.5223 (1969), pp. 957–964.

[35] Gulnara Zh Yusupova et al. “The path of messenger RNA through the ribosome”. In: Cell 106.2 (2001), pp. 233–241.

[36] Aditya Radhakrishnan et al. “The DEAD-box protein Dhh1p couples mRNA decay and translation by monitoring codon optimality”. In: Cell 167.1 (2016), pp. 122–132.

[37] Daniel R Schoenberg. “Mechanisms of endonuclease-mediated mRNA decay”. In: Wiley Interdisciplinary Reviews: RNA 2.4 (2011), pp. 582–600.

[38] Gunter Meister and Thomas Tuschl. “Mechanisms of gene silencing by double-stranded RNA”. In: Nature 431.7006 (2004), pp. 343–349.

[39] Marco Antonio Valencia-Sanchez et al. “Control of translation and mRNA degradation by miRNAs and siRNAs”. In: Genes & development 20.5 (2006), pp. 515–524.

[40] David P Bartel. “MicroRNAs: genomics, biogenesis, mechanism, and function”. In: cell 116.2 (2004), pp. 281–297.

[41] Hermioni Zouridis and Vassily Hatzimanikatis. “A model for protein translation: polysome self-organization leads to maximum protein synthesis rates”. In: Biophysical journal 92.3 (2007), pp. 717– 730.

[42] Atilio Deana and Joel G Belasco. “Lost in translation: the influence of ribosomes on bacterial mRNA decay”. In: Genes & development 19.21 (2005), pp. 2526–2533.

[43] Sergio Barberan-Soler, Nicole J Lambert, and Alan M Zahler. “Global analysis of alternative splicing uncovers developmental regulation of nonsense-mediated decay in C. elegans”. In: Rna 15.9 (2009), pp. 1652–1660.

[44] Erin E Heyer and Melissa J Moore. “Redefining the translational status of 80S monosomes”. In: Cell 164.4 (2016), pp. 757–769.

[45] Stephen N Floor and Jennifer A Doudna. “Tunable protein synthesis by transcript isoforms in human cells”. In: Elife 5 (2016), e10921.

[46] Elodie Zhang et al. “A specialised SKI complex assists the cytoplasmic RNA exosome in the absence of direct association with ribosomes”. In: The EMBO journal 38.14 (2019), e100640.

[47] Rafal Tomecki and Karolina Drazkowska. “An integrative approach uncovers transcriptome-wide determinants of mRNA stability regulation in Saccharomyces cerevisiae”. In: The FEBS Journal 288.11 (2021), pp. 3418–3423.

