## Supplementary for "Factors regulating translation coupled mRNA degradation in yeast"

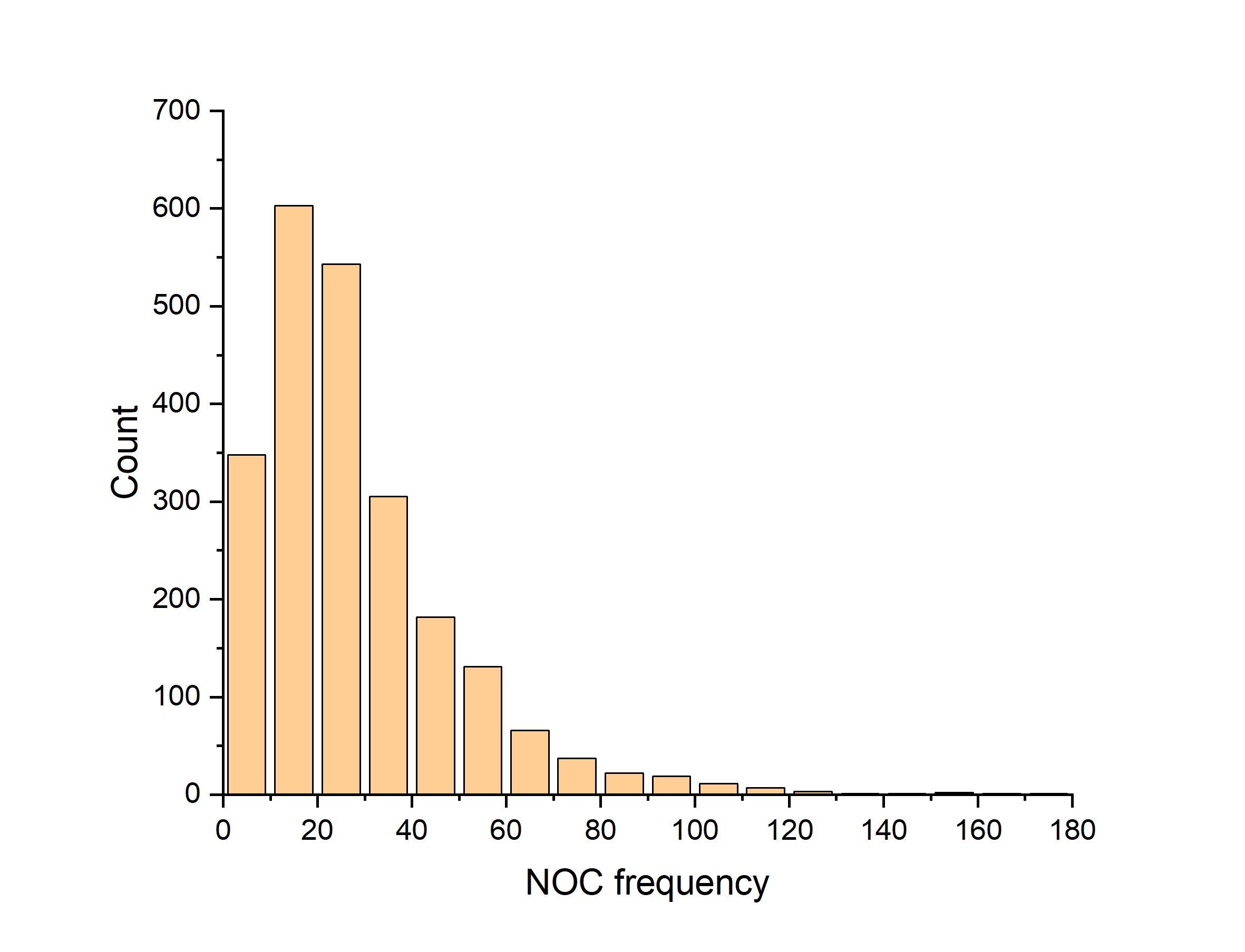
**Fig S1A.** Histogram showing the frequency of non-optimal codon (NOC) stretch (≥3 consecutive non-optimal codon) along 2283 transcripts.

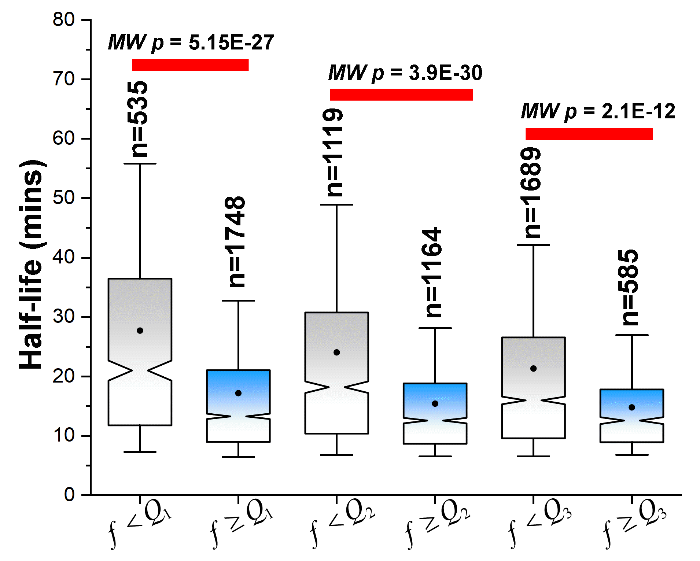
**Fig S1B.** Comparative notched box-plot showing the effect of frequency of NOC stretches on the transcript’s half-life. The NOC stretch distribution is divided into three quartiles (*Q_1_*, *Q_2_* and *Q_3_*). The comparative notched box-plot here represents transcripts belonging (i)below (grey) and (ii) equal and above(blue) a quartile range. Transcripts belonging of lower frequency group are more stable compared to higher frequency groups. The distributions are tested pairwise using two-sample Mann-Whitney test for median and the p-values are provided.

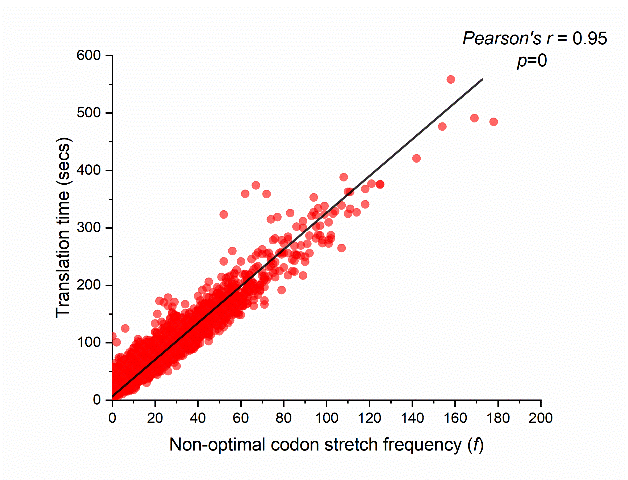
**Fig S1C**. Scatter plot depicting the correlation between Translation time (seconds) (Data S1) and non-optimal codon stretch frequency (*f*). As observed, increase in stretches of non-optimal codon increases the translational time of the transcript (Pearson’s *r*= 0.95, *p*=0), incurring into frequent stoppages.

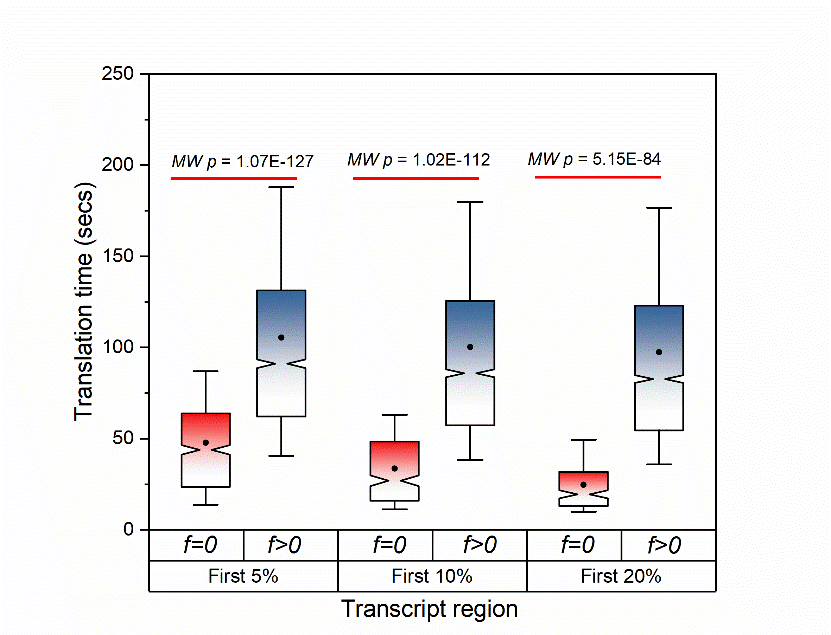

**Fig S1D.** Effect of the absence and presence of non-optimal codon stretch (≥3 stretch length) on the translation time (Data S1). Presence of non-optimal codon stretch at the beginning of the transcript, elevates its translation time.

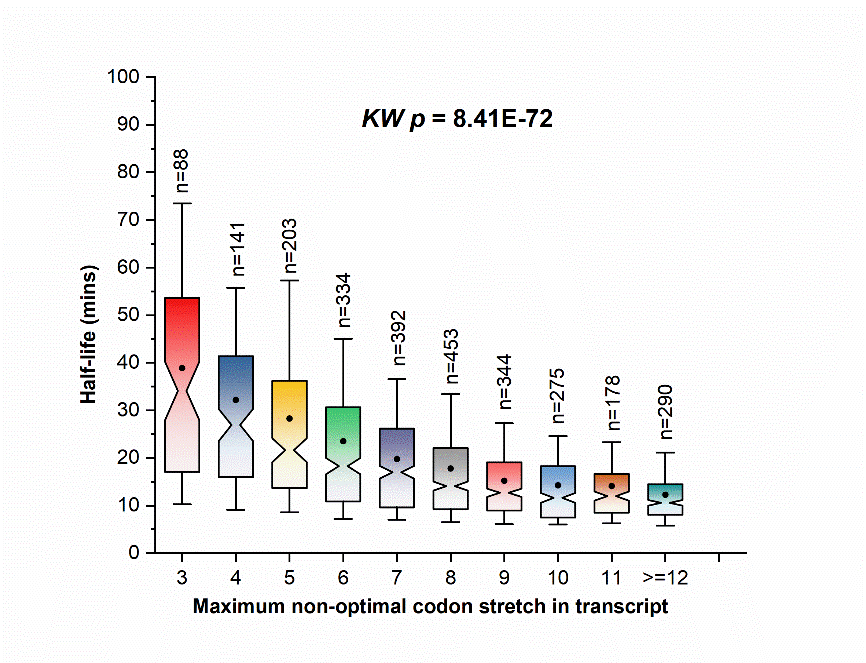
**Fig S1E**. Notched box plot showing the effect of maximum non-optimal codon stretch (NOC) on the transcript half-life. The half-life of the transcript gradually decreases with increase in the maximum NOC stretch within the transcript. The distributions are tested using multi-sample Kruskal-Wallis and the p-value is provided.

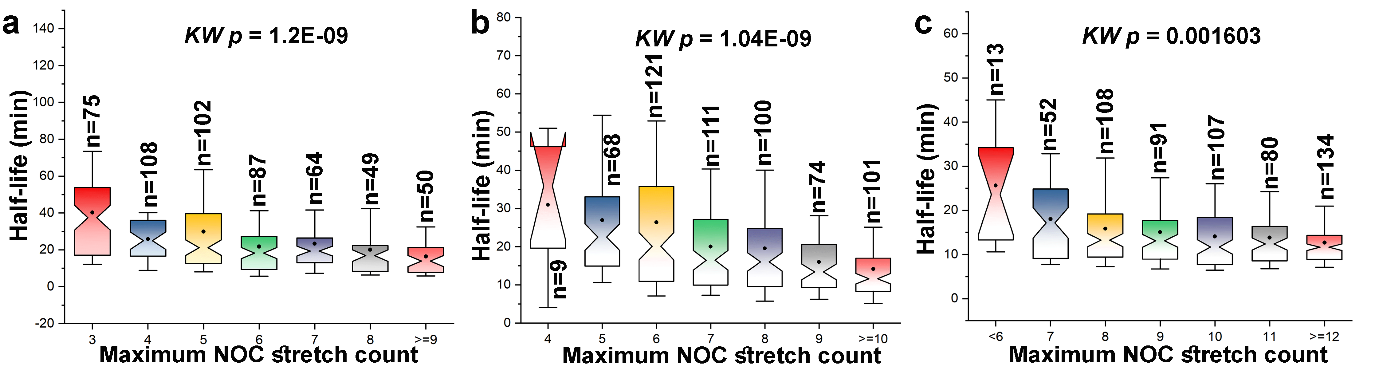
**Fig S1F.** Transcripts belong in each NOC stretch frequency group (a) *f < Q_1_*, (b) *Q_1_ ≤ f < Q_2_*, and (c)*f ≥ Q_3_*, are further classified based on the presence of maximum NOC stretch length. For each frequency group, transcript’s stability gradually decreases with increase in the length of maximum NOC stretch. The distributions for each group are tested using Multi-sample Kruskal Wallis test for medians and the p-values are provided.

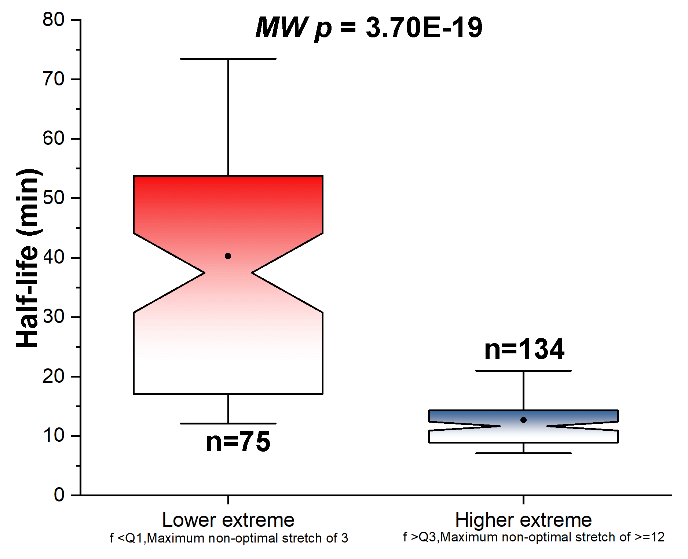

**Fig S1G.** Comparison of transcript’s half-life belonging to (i) Lower extreme, with NOC stretch frequency(*f*) less than *Q_1_* and a maximum NOC stretch length of 3 and (ii) Higher extreme, with NOC stretch frequency(*f*) more than *Q_3_* and a maximum NOC stretch length of above 11. The difference in half-life is tested using pairwise Mann-Whitney test and the p-value is provided.

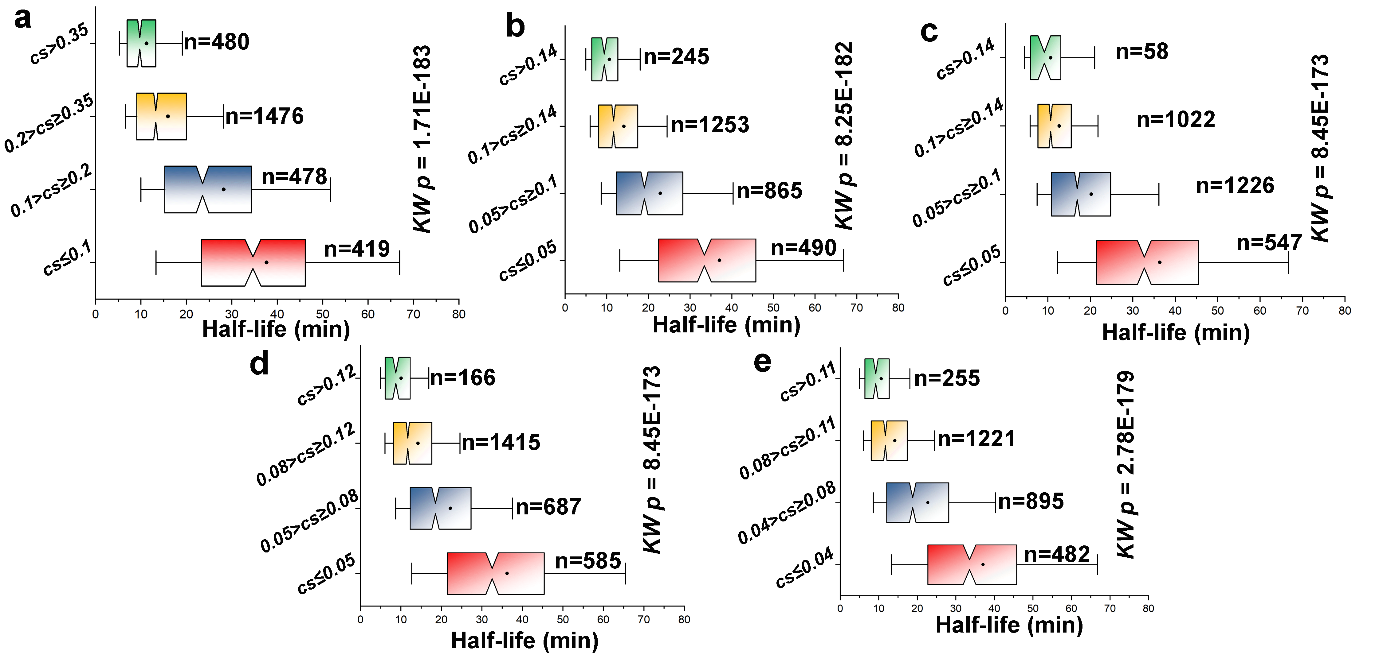
**Fig S1H.** Cumulative score (*cs*) is a unified metric to consider both the frequency and length of non-optimal codon stretch. The metric *cs* includes a weightage factor *k* which is varied, from 0.1 to 10. Increasing the value of *k*, decreases the contribution of frequency, and increases the contribution of maximum NOC stretch length on *cs*. The plot for *k=2* is shown in main text (Fig 1F) while for *k* (a) 1, (b) 3, (c) 4, (d) 5 and (e) 6 is shown here. The stability of transcripts belonging to higher *cs-*group is less that transcripts belonging to lower *cs*-groups. Multi-sample Kruskal Wallis (*KW*) test is performed and the p-value is provided to test whether the distributions differ significantly. A table for all the *KW* p-value for all the values of *k* is given in Table S2.

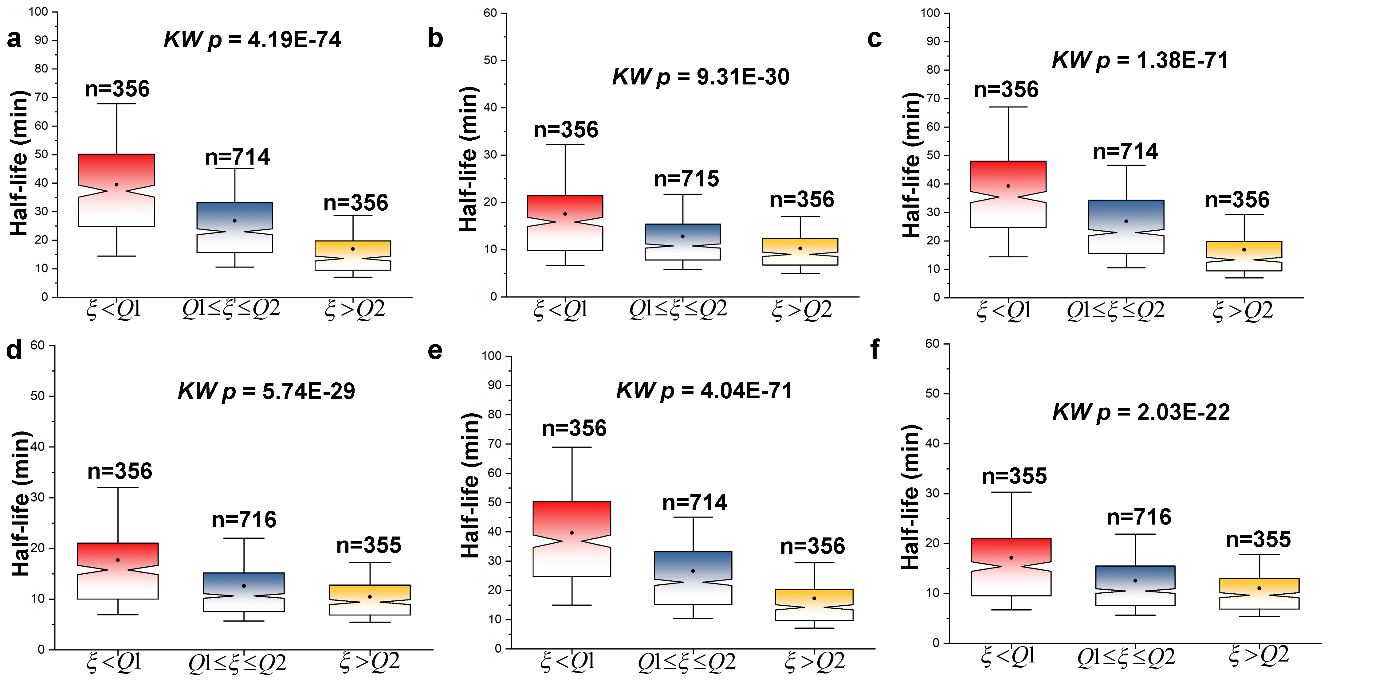
**Fig S2A.** Transcripts are classified based on median (M) *cs* score into two groups. Transcripts belonging to each *cs*-group are further classified based on their *IUS(*
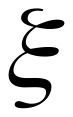
value). *IUS (*
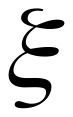
) classification are based on two quartile regions. Half-lives are plotted for *IUS* calculated at structured threshold, (a) and (b) ≥30% for *cs < M* and *cs ≥ M* respectively, (c) and (d) ≥40% for *cs < M* and *cs ≥ M* respectively, (e) and (f) ≥70% for *cs < M* and *cs ≥ M* respectively. The plot for half-life for IUS threshold for structured threshold ≥60% for *cs < M* and *cs ≥ M* is given main text in Fig 2A and 2B.

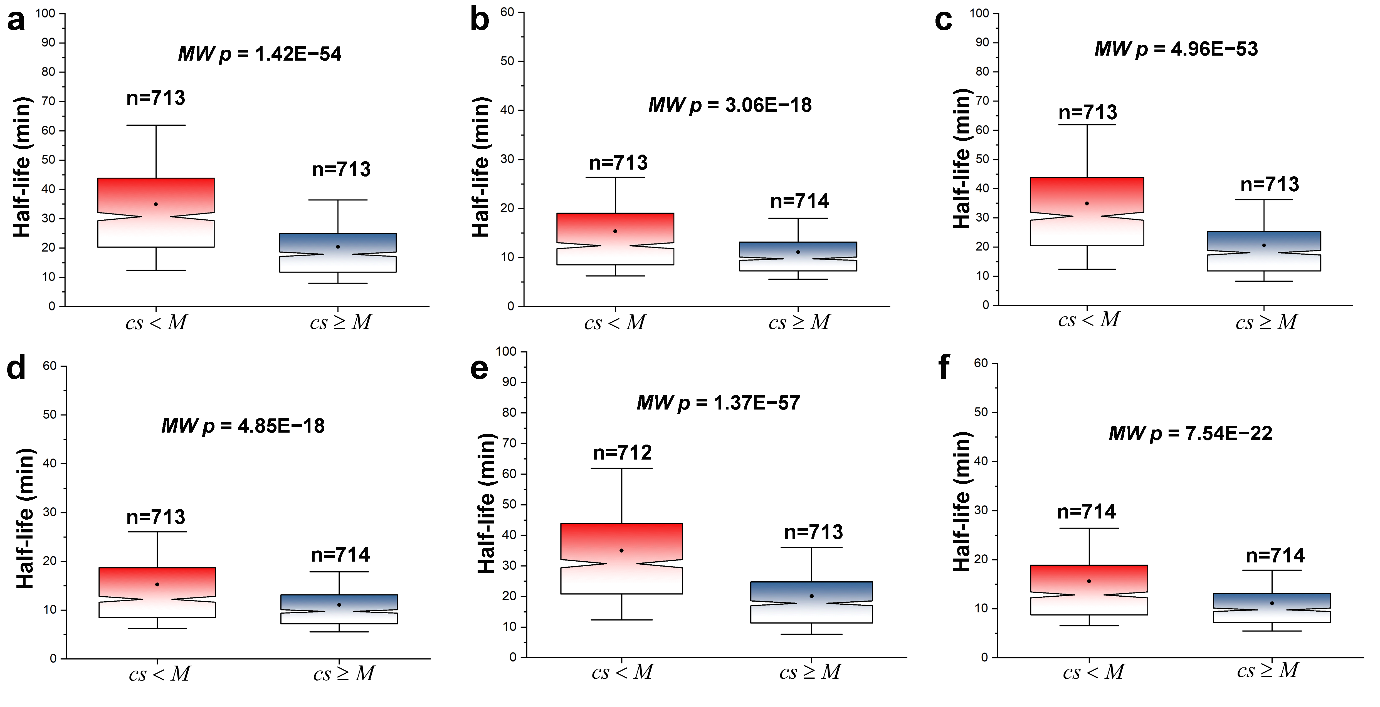
**Fig S2B.** Notched box plot representing half-life of transcript which are initially classified based in median *IUS(*
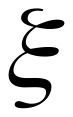
) into two groups (
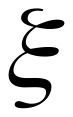
 < M and
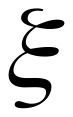
 ≥ M). The transcripts in each group are further classified based on the median *cs*-score. Half-life plotted for transcript’s
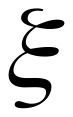
 at structured threshold of, (a) and (b) ≥30% for
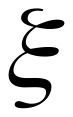
 < M and
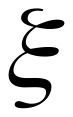
 ≥ M respectively, (c) and (d) ≥40% for
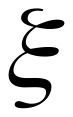
 < M and
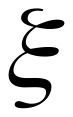
 ≥ M respectively, (e) and (f) ≥70% for
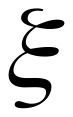
 < M and
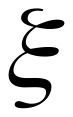
 ≥ M respectively. The plot for half-life for IUS threshold for structured threshold ≥60% for
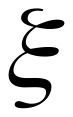
 < M and
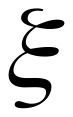
 ≥ M is given main text in Fig 2C and 2D.

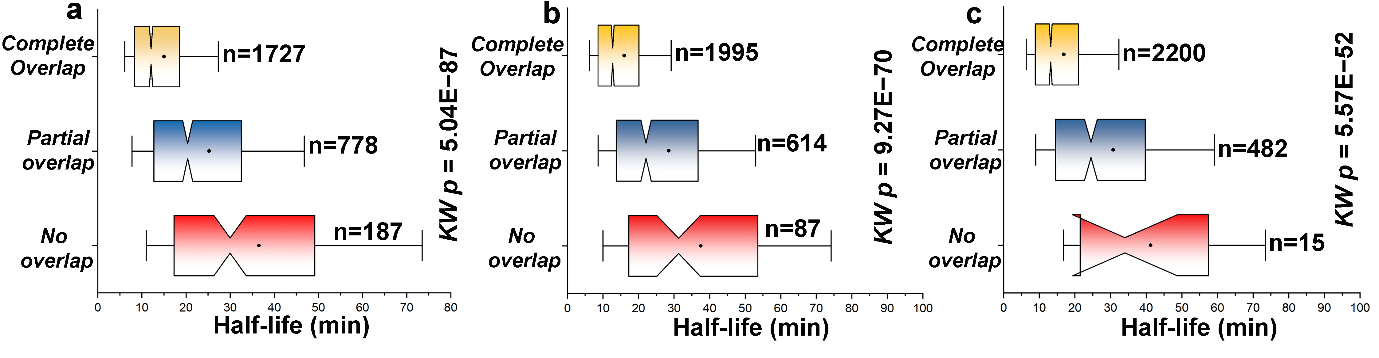
 **Fig S2C.** Notched box plot of half-life for transcripts having complete, partial and no overlap between *IUS* regions and non-optimal codon stretches. The plots are generated for
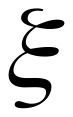
 values with structured threshold of (a) ≥30%, (b) ≥40% and (c)≥70%. The plot for overlap between non-optimal codon stretch and *IUS* for structural threshold ≥60% is shown in main text in Fig 2E. Transcripts having unstructured regions with the NOC stretches or even in vicinity have reduced stability, as unstructured segments provide amenable sites for nuclease attachment and degradation after transcript release.

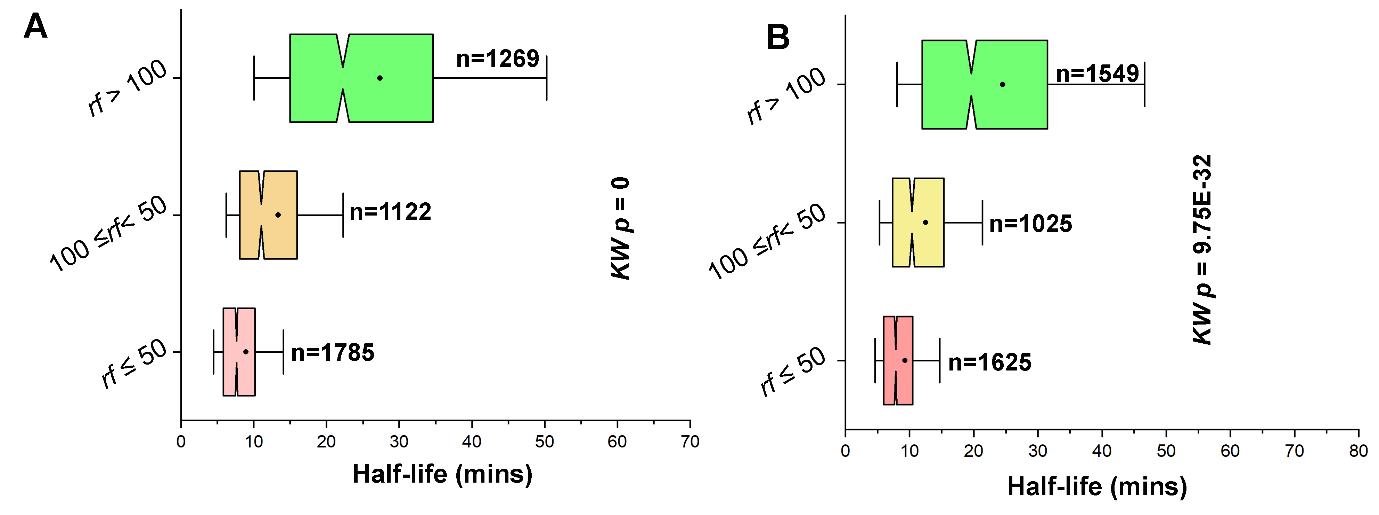
 **Fig S3A.** Effect of ribosomal footprinting (*rf)* on transcript’s half-life. There are three *rf* datasets, which is based on the data extraction method, which are Ribominus, Unselected and Dynabeads. Here, we have notched box plot for (A) Unselected and (B) Dynabeads, while the plot for Ribominus is given in main text. We see increase in sequestration of mRNA with ribososme elevates stability and hence, increase in half-life is observed.

**Fig S3B.** Transcripts are initially classified based in the median *cs-*score into two groups (*cs* < M and *cs*≥M). Each of this group is further divided based on their *rf* value. The plots here are shown for (a) and (b)
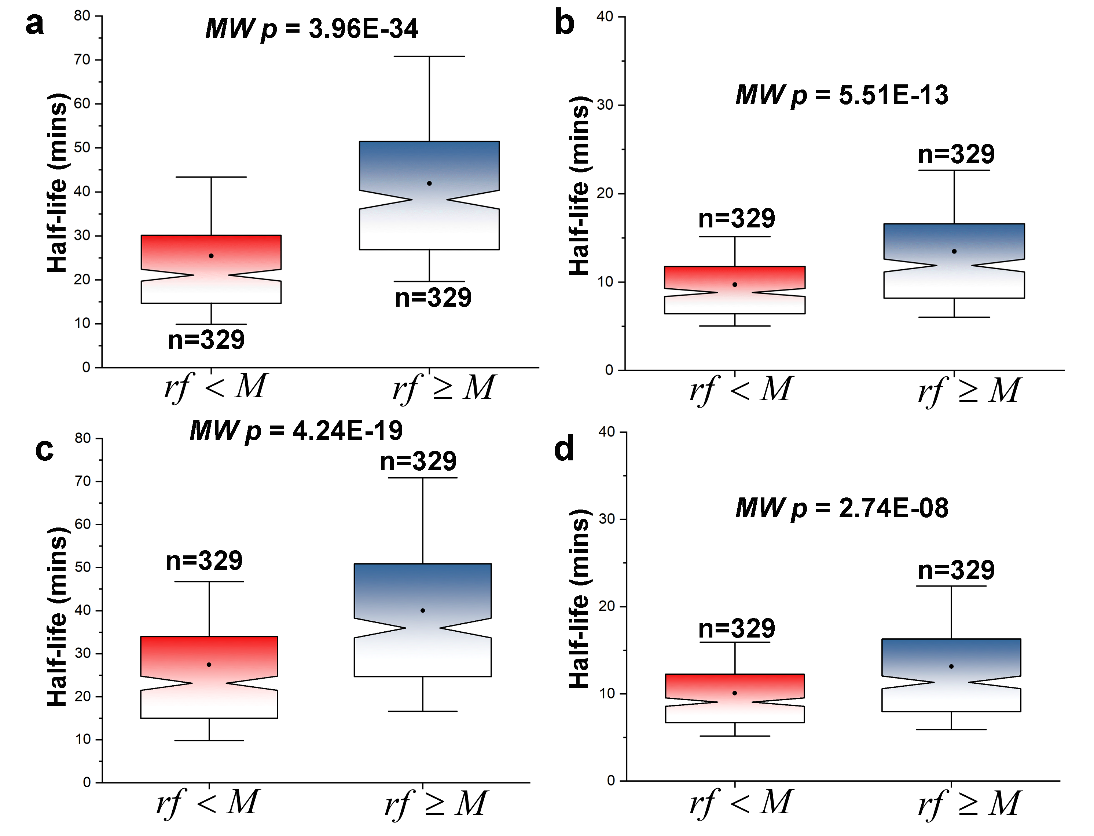
Unselected with *cs<M* and *cs≥M* respectively, and (c) and (d) Dynabeads with *cs<M* and *cs≥M* respectively.

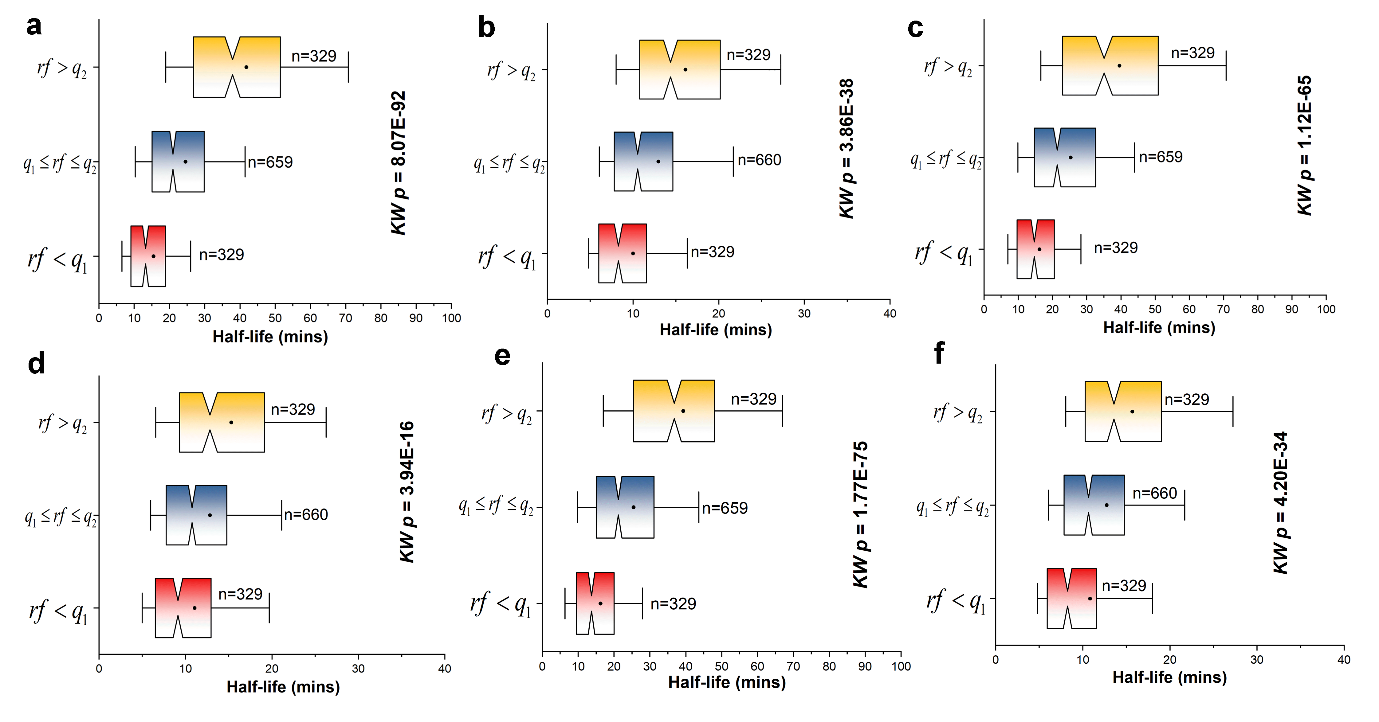

**Fig S3C.** Transcripts are initially classified based on *IUS (*
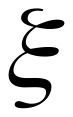
) at structured threshold of ≥30% into two groups,
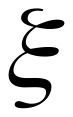
<M and
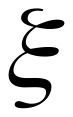
≥M. Notched box plot representing half-life of transcript for different range of *rf* values for three datasets, (a) and (b) Unselected for
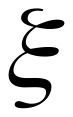
<M and
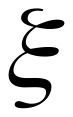
≥M respectively, (c) and (d) Dynabeads for
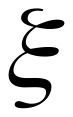
<M and
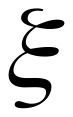
≥M respectively, and (e) and (f) Ribominus for
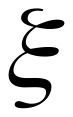
<M and
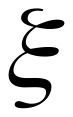
≥M respectively.

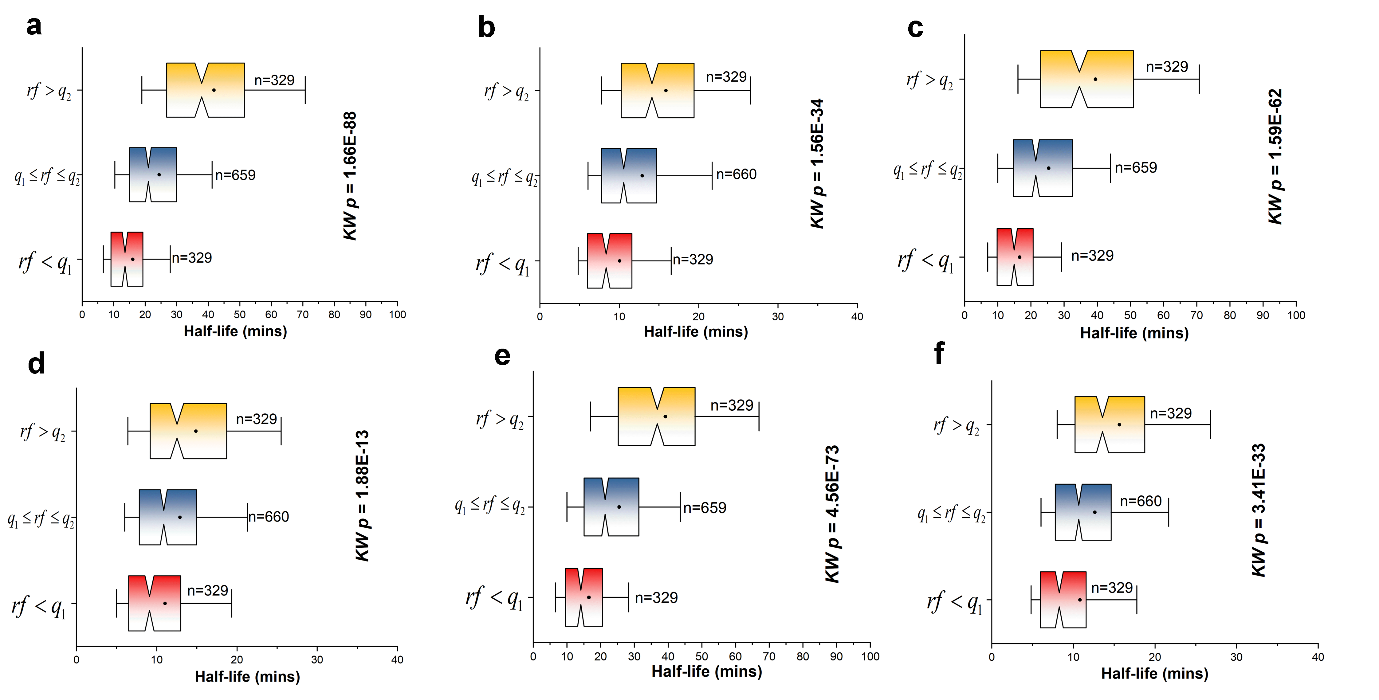

**Fig S3D.** Transcripts are initially classified based on *IUS (*
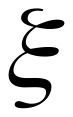
) at structured threshold of ≥40% into two groups,
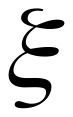
<M and
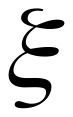
≥M. Notched box plot representing half-life of transcript for different range of *rf* values for three datasets, (a) and (b) Unselected for
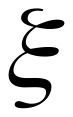
<M and
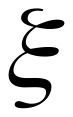
≥M respectively, (c) and (d) Dynabeads for
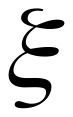
<M and
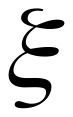
≥M respectively, and (e) and (f) Ribominus for
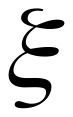
<M and
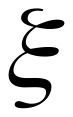
≥M respectively.

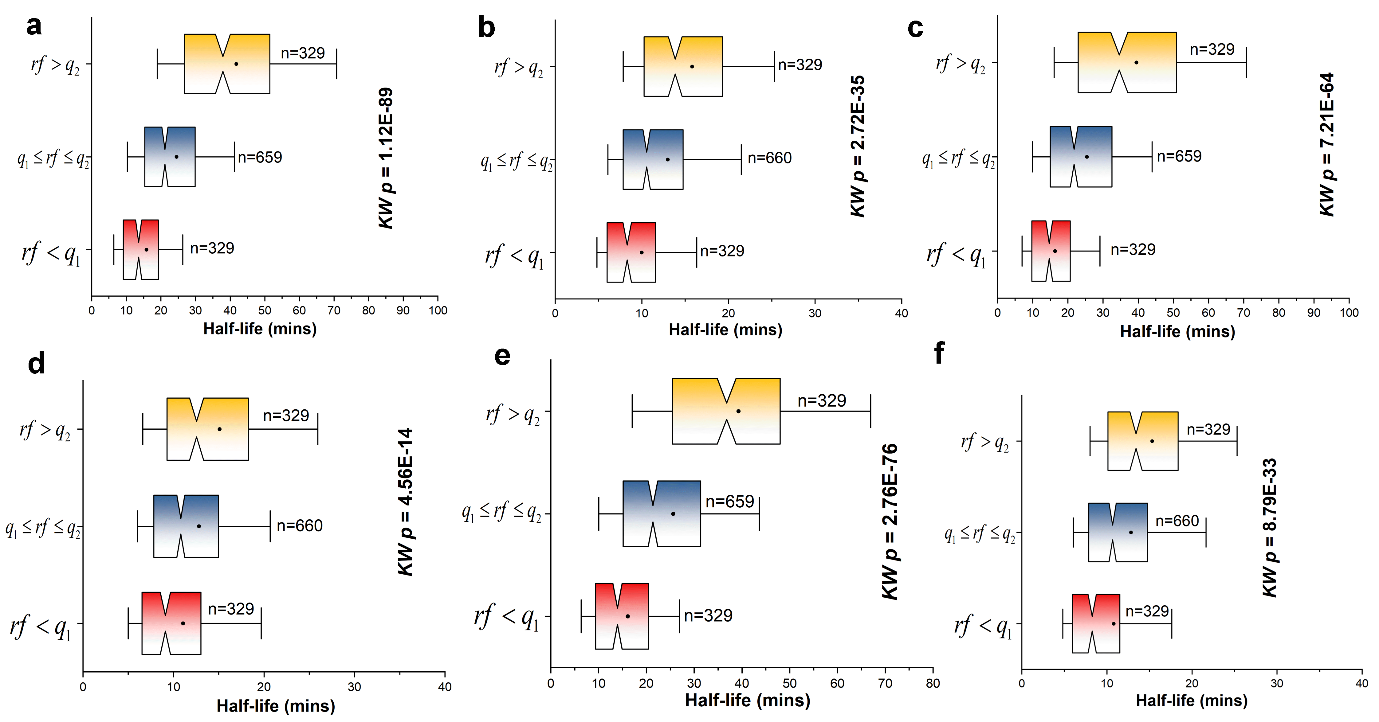
**Fig S3E.** Transcripts are initially classified based on *IUS (*
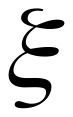
) at structured threshold of ≥60% into two groups,
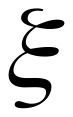
<M and
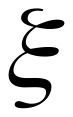
≥M. Notched box plot representing half-life of transcript for different range of *rf* values for three datasets, (a) and (b) Unselected for
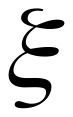
<M and
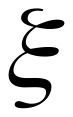
≥M respectively, (c) and (d) Dynabeads for
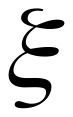
<M and
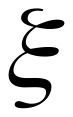
≥M respectively, and (e) and (f) Ribominus for
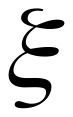
<M and
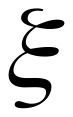
≥M respectively.

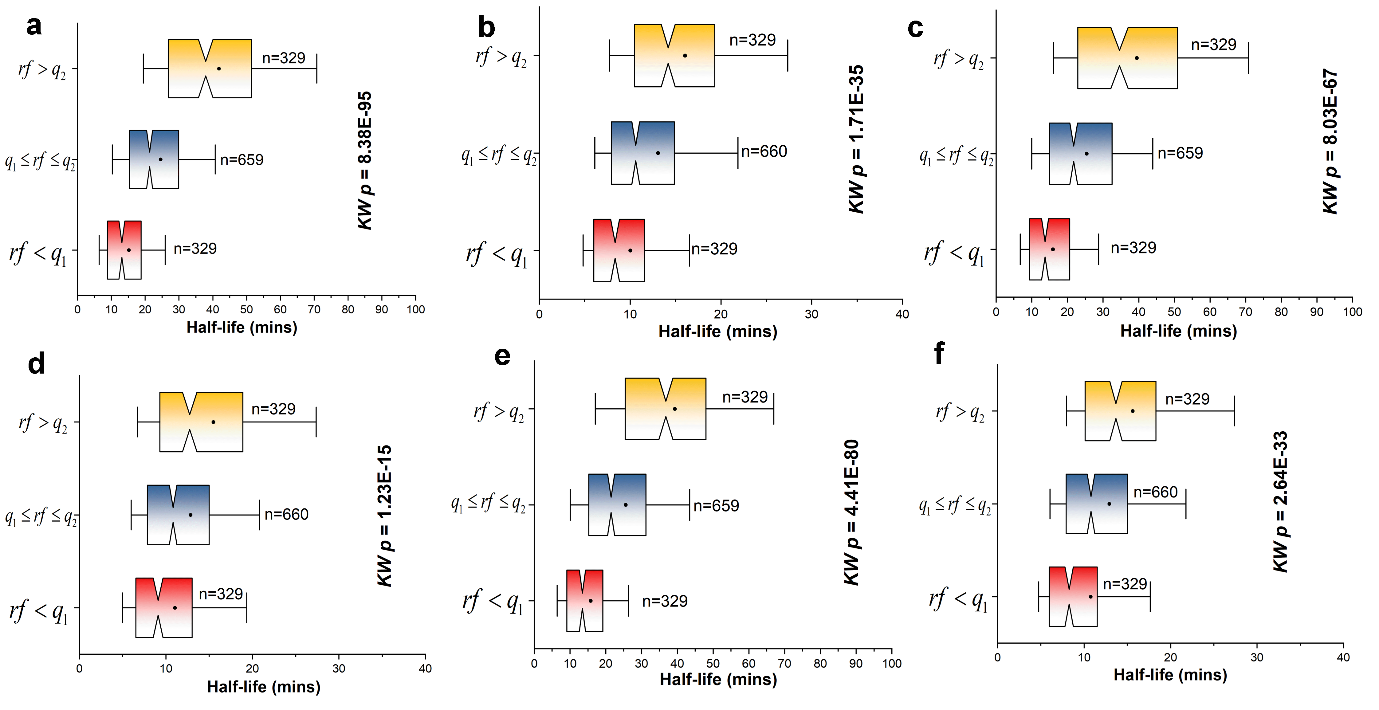

**Fig S3F.** Transcripts are initially classified based on *IUS (*
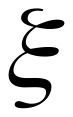
) at structured threshold of ≥70% into two groups,
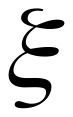
<M and
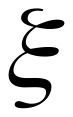
≥M. Notched box plot representing half-life of transcript for different range of *rf* values for three datasets, (a) and (b) Unselected for
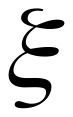
<M and
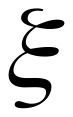
≥M respectively, (c) and (d) Dynabeads for
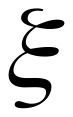
<M and
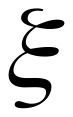
≥M respectively, and (e) and (f) Ribominus for
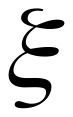
<M and
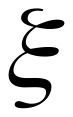
≥M respectively.

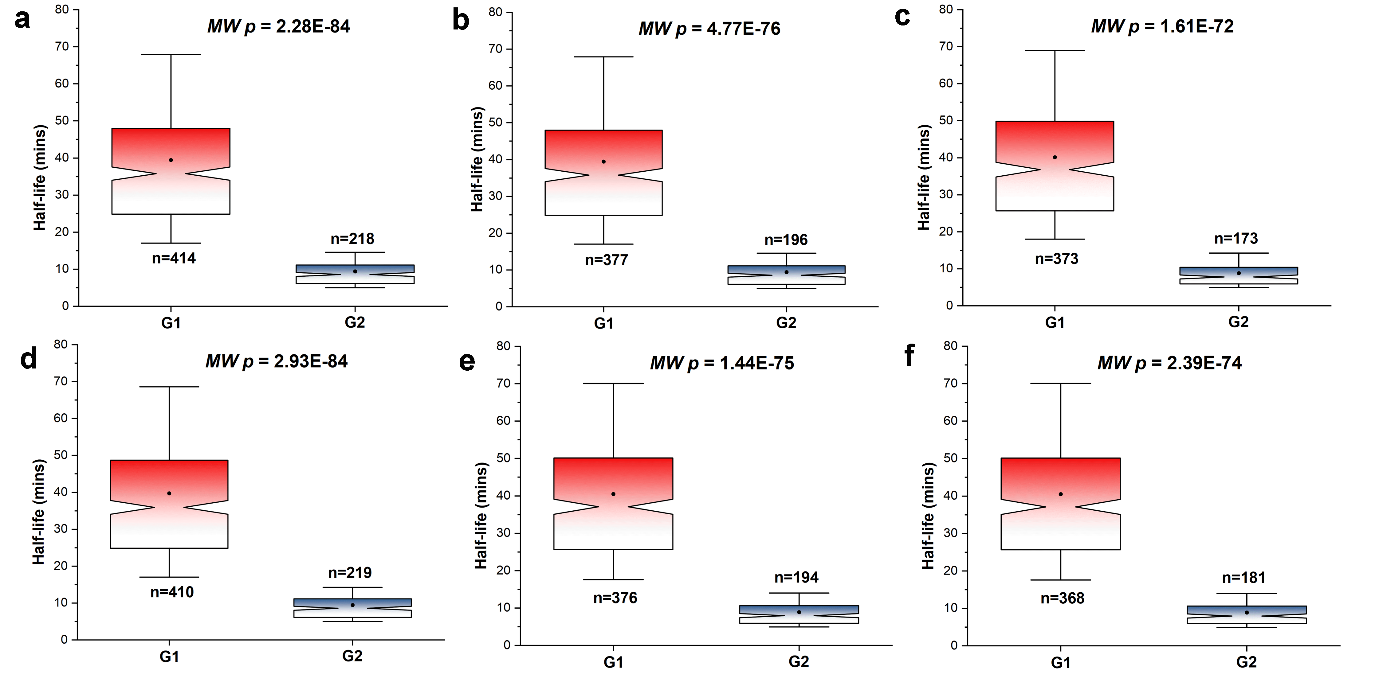
**Fig S3G.** Transcripts are classified based on two extreme conditions, (i)
cs < Q1, ξ < Q1 and rf > Q3 referred as G1 and (ii) cs > Q3, ξ > Q3 and rf < Q1 referred as G2.
The half-life between these two extremes is compared. The plot here are calculated for *IUS* structured threshold of (a), (b) and (c) ≥30% for Unselected, Dynabeads and Ribominus respectively and (d), (e) and (f) ≥40% for Unselected, Dynabeads and Ribominus respectively.

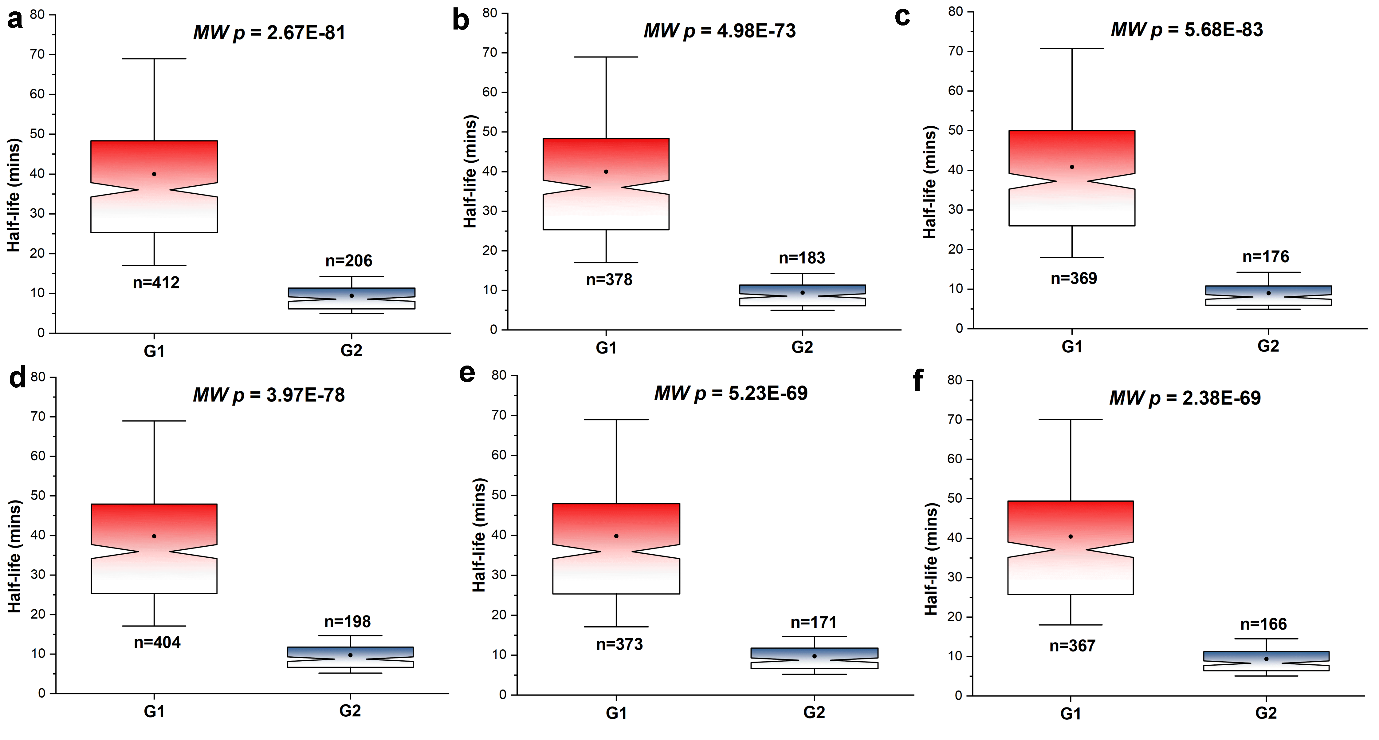

**Fig S3H.** Transcripts are classified based on two extreme conditions, (i)
cs < Q1, ξ < Q1 and rf > Q3 referred as G1 and (ii) cs > Q3, ξ > Q3 and rf < Q1 referred as G2.
The half-life between these two extremes is compared. The plot here are calculated for *IUS* structured threshold of (a), (b) and (c) ≥60% for Unselected, Dynabeads and Ribominus respectively and (d), (e) and (f) ≥70% for Unselected, Dynabeads and Ribominus respectively.

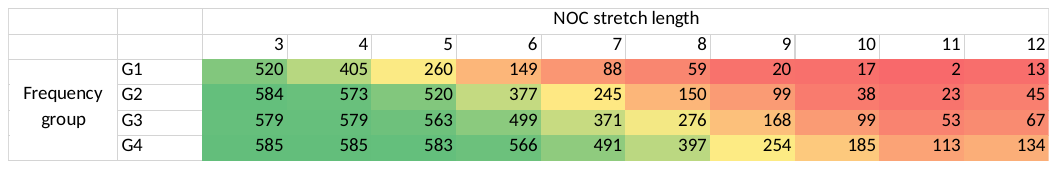

**Table S1**. Table showing the distribution of different *NOC* stretch length (≥3) within the transcript belonging to each frequency group. Lower *NOC* stretches are common among all frequency groups, while higher *NOC* stretch length (≥5) significantly increase among transcripts belonging to higher frequency group.

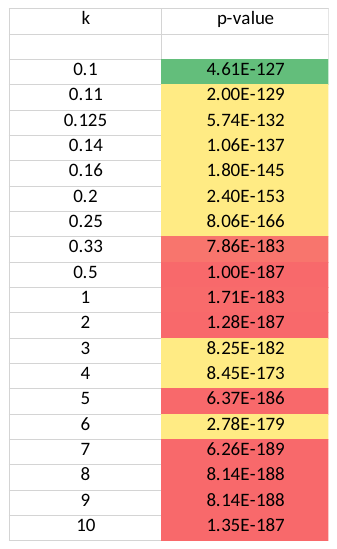

**Table S2.** A weightage factor (*k*) is used in the metric of *cs*. The weightage factor is varied from 0.1 to 10, in order to differential effect of both frequency and non-optimal codon stretch length onto the *cs*-score. For *k<1,* the effect of frequency dominates, while for *k>1,* the non-optimal codon stretch dominates on *cs.* Half-life’s of transcripts is plotted as notched box plot for different range of *cs-*value at different value of *k.* The distributions are statistically tested using multi-sample Kruskal-wallis test of medians. The *p-values­* for those distribution plot at different *k* value is shown in this table.
